# Theoretical assessment of persistence and adaptation in weeds with complex life cycles

**DOI:** 10.1101/2022.08.12.503772

**Authors:** Dana Lauenroth, Chaitanya S. Gokhale

## Abstract

Herbicide-resistant weeds pose a significant threat to global food security. Perennial weed species are particularly troublesome. Such perennials as *Sorghum halepense* spread quickly and are difficult to manage due to their ability to reproduce sexually via seeds and asexually through rhizomes. Our theoretical study of *Sorghum halepense* incorporates this complex life cycle with control measures of herbicide application and tillage. Rooted in the biology and experimental data of *Sorghum halepense*, our population-based model predicts population dynamics and target-site resistance evolution in this perennial weed. We found that the resistance cost determines the standing genetic variation for herbicide resistance. The sexual phase of the life cycle, including self-pollination and seed bank dynamics, contributes substantially to the persistence and rapid adaptation of *Sorghum halepense*. While self-pollination accelerates target-site resistance evolution, seed banks significantly increase the probability of escape from control strategies and maintain genetic variation. Combining tillage and herbicide application effectively reduces weed densities and the risk of control failure without delaying resistance adaptation. We also show how mixtures of different herbicide classes are superior to rotations and monotreatment in controlling perennial weeds and resistance evolution. Thus, by integrating experimental data and agronomic views, our theoretical study synergistically contributes to understanding and tackling the global threat to food security from resistant weeds.

## Introduction

*Sorghum halepense* (Johnsongrass) is infamous for being “a threat to native grasslands and agriculture” (Klein & Smith 2021, p. 413). Johnsongrass is classified as a weed in 53 countries (Klein & Smith 2021), and Holm et al. (1977) ranked it as the world’s sixth-worst weed. It causes significant yield losses in 30 different crops, including essential cash crops such as corn, cotton, sugarcane and soybean (Klein & Smith 2021). The grassy weed out-competes native grasses like big bluestem (*Andropogon gerardii*), little bluestem (*Schizachyrium scoparius*) and switchgrass (*Panicum virgatum*) in grasslands (Schwinning et al. 2017). Johnsongrass is a highly competitive and invasive perennial, capable of sexual reproduction via seeds and asexual propagation through rhizomes (Warwick & Black 1983, Peerzada et al. 2017, Schwinning et al. 2017). In addition to resource competition, Johnsongrass affects crop production by serving as an alternative host for various agricultural pests (Peerzada et al. 2017, Klein & Smith 2021).

In cropped areas, typical control of Johnsongrass is by a post-emergence application of herbicides, particularly glyphosate, acetyl-coenzyme-A carboxylase (ACCase)- and acetolactate synthase (ALS)-inhibitors (Peerzada et al. 2017, Klein & Smith 2021). However, the continuous use of such herbicides has caused Johnsongrass to develop resistances, including target-site resistance towards various ACCase- and ALS-inhibitors (Lorentz 2014, Peerzada et al. 2017) and non-target-site resistance to glyphosate (Vila-Aiub et al. 2007). Herbicide resistance imposes a considerable challenge on weed management. There is an urgent need for weed management strategies that reduce herbicide resistance risk while effectively controlling the weed population. Integrated approaches, combining herbicide application with mechanical measures, such as tillage, and potentially crop rotations, are reported to provide efficient control of Johnsongrass, including its rhizomes (Warwick & Black 1983, Peerzada et al. 2017). Mixtures and rotations of herbicides with different sites of action reportedly reduce weed densities more efficiently than recurrent use of one herbicide class and delay resistance evolution (Diggle et al. 2003, Busi et al. 2020, Zeller et al. 2021).

To develop and select sustainable control strategies, we must improve our understanding of eco-evolutionary processes underlying the population dynamics and how inherent plant characteristics shape the evolution of herbicide resistance in perennial weeds. Mathematical modelling has proven valuable in studying long-term population dynamics and herbicide resistance evolution in weeds (Holst et al. 2007, Freckleton & Stephens 2009). However, the majority of existing models deal with annual weeds; the complex life cycle of perennial species has been addressed to a lesser extent (Holst et al., 2007, but see Liu et al., 2019; Holmes et al., 2022). To our knowledge, only two population dynamical models exist for Johnsongrass, and more general for perennials comprising the whole life cycle (Liu et al. 2019, Holmes et al. 2022). Liu et al. (2019) presented an individual-based model, which captures self-pollination but lacks a seed bank. The stage-structured model of Holmes et al. (2022) comprises both a seed bank and a bank of rhizome buds. However, the model lacks the possibility of de novo resistance evolution, and pure cross-pollination is assumed.

The goals of this study are (i) to explore how the resistance cost and its dominance affect the standing genetic variation for target-site resistance; (ii) to study how genetic details, like the dominance of resistance and the associated fitness cost, impact population dynamics and target-site resistance evolution in herbicide-treated weeds; (iii) to examine the effects of self-pollination and seed bank formation on genetic diversity, target-site resistance evolution, and risk of control failure in perennial species; (iv) to determine the potential of tillage, herbicide mixtures and rotations for controlling resistance evolution and weed control failure. To this aim, we implemented the life cycle of Johnsongrass, both mathematically and computationally. The sexual and asexual phases are incorporated differentially since the spread of resistance is explicitly genetic.

Even though the model is presented in the context of Johnsongrass, it can capture the population dynamics and herbicide resistance adaptation of general perennial weeds. More generally, our population-based approach can be adapted to any species that comprise sexual reproduction in the continuum of selfing and outcrossing, asexual propagation, and potentially a seed bank component.

## Model summary

The seminal work by Sager & Mortimer (1976) has inspired several models of weed life cycles. The population dynamic is typically captured deterministically using several discrete-time equations, each corresponding to the included life stages (Holst et al. 2007). Often the population is assumed to be even-aged, with the system time-steps of one-year increments (Holst et al. 2007). We introduce a detailed population-based model describing the dynamics of seed- and rhizome-propagated Johnsongrass. Our deterministic model with an annual time step comprises the whole perennial life cycle. We then use the model to forecast population dynamics and the evolution of target-site resistance. In this section, we summarise the main features of our theoretical model: life history stages and eco-biology, control measures and resistance, and stochastic simulations. For a detailed exposition, we refer to the Methods. We inferred all model parameters carefully from field trials in the literature. In the Supplementary Information, we provide Table 1 with all parameter values and the derivation.

### Life cycle

Johnsongrass is dormant throughout the winter, overwintering as seeds or rhizomes in the ground (Warwick & Black 1983). At the beginning of a growing season, seeds germinate to produce seedlings, and shoots emerge from nodes on the rhizomes (Peerzada et al. 2017). Mature plants produce new rhizomes and, after flowering, viable seeds (Warwick & Black 1983, Peerzada et al. 2017).

Figure 1 outlines the life cycle of Johnsongrass with both reproduction pathways, sexual via seeds and asexual through rhizomes (a) and its representation in our model (b). The figure further illustrates the implementation of self-thinning and density-dependent reproduction resulting from intraspecific competition (c). Only rhizomes and seeds are present in early spring. Axillary buds formed on the nodes of rhizomes can develop as shoots or secondary rhizomes (Paterson et al. 2020). However, the apex of a rhizome exerts apical dominance over axillary buds, inhibiting their growth (Beasley 1970). The number of shoots in any given season can be determined from the number of rhizomes, the number of nodes on rhizomes and the proportion of nodes producing shoots.

**Figure 1:**
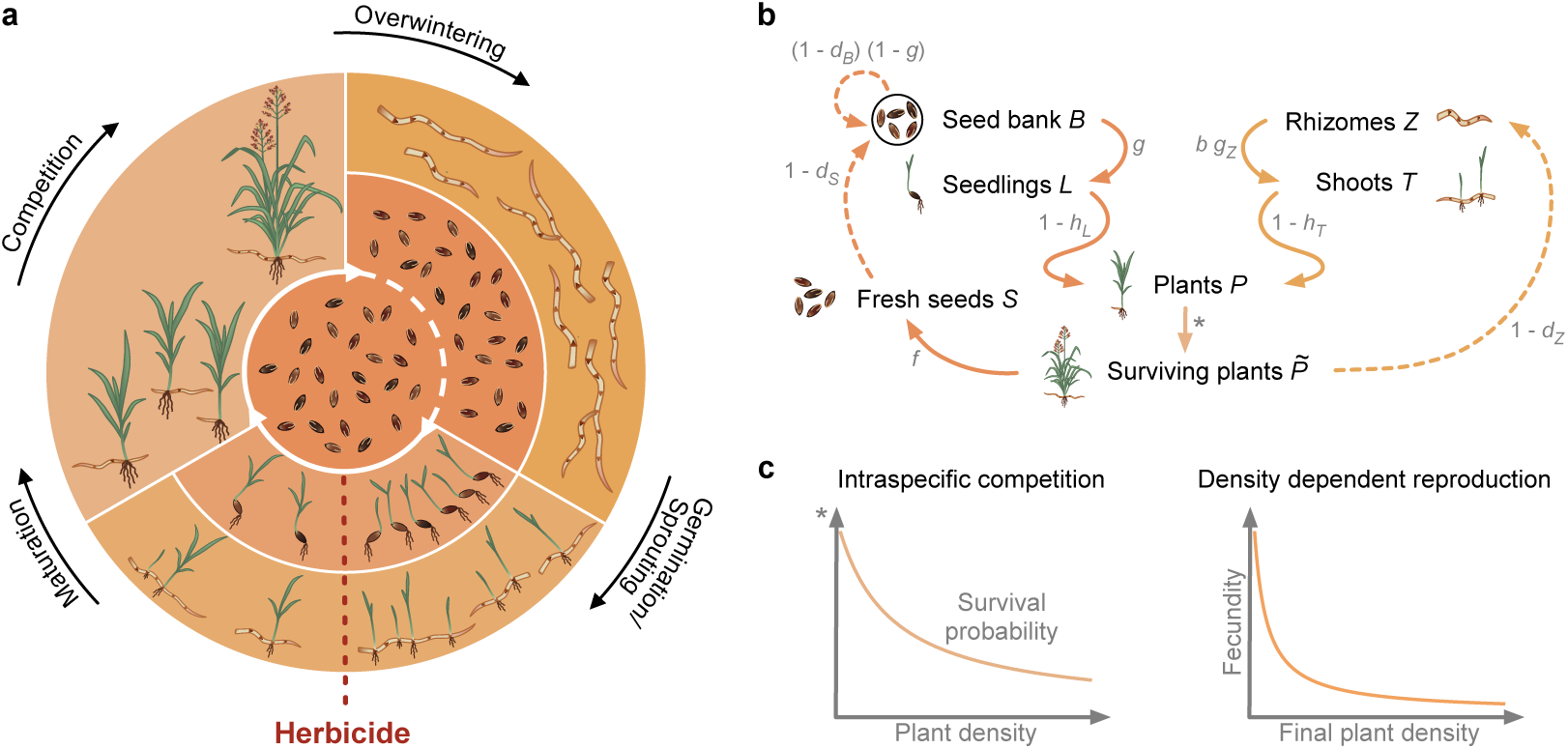
Schematic illustration of Johnsongrass’ life cycle and its representation in our model. **a**, Diagram of Johnsongrass’ life cycle. Johnsongrass reproduces sexually via seeds (inner ring) and asexually through rhizomes (outer ring). Seeds can stay dormant in the ground for several years, forming a seed bank (central circle). New seeds and seeds from the seed bank might germinate in spring or stay dormant as part of the seed bank (expressed by the dotted line). Rhizomes give rise to shoots in the first spring after their production. Herbicide application (red dotted line) can kill susceptible seedlings and shoots. The plants that survive then compete for resources as they mature. The aboveground plant material dies in winter, and Johnsongrass overwinters as seeds and rhizomes in the ground. **b**, Schematic representation of our model. The left part of (b) corresponds to the sexual reproduction of Johnsongrass, and the right part represents the asexual propagation. Solid arrows depict within-season dynamics, and dashed arrows show dynamics between seasons. Survival probabilities and fecundity are shown in grey next to the corresponding arrows. **c**, Intraspecific competition. Intraspecific resource competition leads to self-thinning and density-dependent fecundity reduction. The left panel displays the probability of intraspecific competition survival in young plants (*P*) as a function of their density. The density-dependent reduction in fecundity, i.e. the number of seeds (*f*) and rhizome buds (*b*) produced by mature plants, is illustrated on the right.

Freshly produced Johnsongrass seeds are highly dormant throughout the season (Warwick & Black 1983). They germinate in the subsequent seasons and can stay viable in the ground for several years (Warwick & Black 1983, Peerzada et al. 2017). New seeds enter the seed bank but are subject to post-dispersal predation and might lose viability or decay until the beginning of the next season (Bagavathiannan & Norsworthy 2013). Each spring, only a certain proportion of seeds in the seed bank germinates to produce seedlings (Legutzamón 1986). Non-germinated seeds may either be lost due to decay and viability loss or stay dormant (Bagavathiannan & Norsworthy 2013). At all times, the seed bank consists of non-germinated seeds from the previous seed bank and seeds produced in the preceding season.

Seedlings and shoots grow into adult plants. Even though rhizome shoots might emerge earlier than seedlings due to a lower temperature requirement for germination, similar growth and development were observed for Johnsongrass originating from seeds and rhizomes (McWhorter 1961, Horowitz 1972). Hence, we do not distinguish the adult plants by their origin. The cycle continues as the adult plants produce new seeds and rhizomes, of which only a certain amount survive the winter.

### Intraspecific competition

Johnsongrass is more strongly affected by intraspecific competition than by competition from other perennial grasses, like big bluestem, little bluestem and switchgrass (Schwinning et al. 2017). In addition, since the crop density is assumed to remain constant throughout the seasons, we do not explicitly model competition from the crop and other weeds. We implement the hyperbolic self-thinning function introduced by Yoda et al. (1963) to model early plant survival (illustrated in Fig. 1 c).

Moreover, Williams & Ingber (1977) reported a significant impact of intraspecific competition on the formation of reproductive structures in Johnsongrass. In their greenhouse experiment, high stand densities delayed or even prevented tillering and delayed the formation of rhizomes and panicles. Also, the final dry weight of reproductive structures was reduced in crowded stands. Due to a lack of data, we assume that the total seed and rhizome yield per square meter approaches a constant value at moderate to high densities (Firbank & Watkinson 1985) (the density-dependent fertility is illustrated in Fig. 1 c).

### Control and resistance

Numerous control strategies exist for managing weed populations. They are typically classified according to their primary mode of action: chemical, mechanical, biological, cultural or allelopathic (Peerzada et al. 2017). We focus on tillage (mechanical control) and postemergence herbicide application (chemical control).

As for target-site resistance, e.g. against ACCase- or ALS-inhibitors, we consider a single allele R associated with herbicide resistance (Scarabel et al. 2014, Hernndez et al. 2015, Kreiner et al. 2017). We assume a diploid genome, but the model can be extended to reflect higher ploidy levels (see Holmes et al. (2022) for a comparison of the rate of herbicide resistance evolution in diploid and tetraploid weeds). The susceptible phenotype results from the homozygous genotype WW, and genotype RR plants are resistant. The factor 0 *≤ k_h_ ≤* 1 captures the variability in the dominance of resistance such that we can account for recessivity (*k_h_*= 0), partial dominance (0 *< k_h_ <* 1) and complete dominance (*k_h_* = 1) of the resistance trait. Spontaneous mutations to the resistance allele R can occur during sexual reproduction. We do not consider back mutations as they do not change the dynamics relevantly. Johnsongrass is primarily self-pollinated, leading to higher homozygosity and reduced genetic diversity in the produced seeds compared to cross-fertilization (Warwick & Black 1983). The sexual reproduction in our model follows Mendelian inheritance and includes spontaneous mutations to the resistance allele R occurring with probability *µ* (see Supplementary Information).

#### Standing genetic variation

Indeed, while mutations can emerge due to the selection pressure of the herbicides, resistance may already exist in the natural variation in a population. Standing genetic variation for herbicide resistance has been found in plant populations never exposed to herbicides, among others, target-site resistance against ACCase-inhibitors (Délye et al. 2013). Moreover, herbicide resistance adaptation may primarily proceed from standing genetic variants as the initial frequency of resistant individuals may be much higher than the spontaneous mutation rate of resistance alleles (Kreiner et al. 2017, Délye et al. 2013). Thus we also include standing genetic variation in our model and derive an analytic approximation of the expected variation (*v*) in an untreated field.

#### Resistance cost

Herbicide resistance can be associated with a fitness cost (Baucom 2019). In particular, a fitness cost on seed production associated with a mutation conferring resistance against ACCase-inhibitors has been reported in Johnsongrass (Panozzo & Sattin 2021). As for the herbicide resistance, the model allows for dominance variability regarding the fitness cost, implemented by a dominance factor 0 *≤ k_c_ ≤* 1. Hence, for type RR and RS plants, a reduction in the average number of seeds produced by *c* and *k_c_c*, respectively, is considered compared to the fecundity of susceptible plants. The resistance cost is always relevant, irrespective of the control regime.

#### Herbicide efficacy

In our model, herbicide resistance is complete for homozygous-resistant plants. Herbicide application can kill susceptible plants and partly resistant heterozygotes with an efficacy *h* and (1 *− k_h_*) *h*, respectively. Due to inadequate translocation of herbicides into the rhizomes and dormant buds, their control is only partial (Beasley 1970, Tuesca et al. 1999). Thus even when the herbicide kills shoots, new shoots may eventually resprout from the parent rhizome (Beasley 1970, Tuesca et al. 1999). This resprouting reduces the herbicide efficacy on rhizome shoots compared to seedlings (Vidrine 1989). Johnsongrass rhizomes are cut into pieces during tillage and partly turned to the soil surface (Warwick & Black 1983). The fragmented rhizomes in shallow layers are exposed to low temperatures during winter, leading to increased mortality (Hull 1970, Warwick & Black 1983). Additionally, rhizome fragmentation can maximize bud sprouting (McWhorter & Hartwig 1965), allowing for enhanced control of Johnsongrass plants originating from rhizomes by the herbicide (Hull 1970, Tuesca et al. 1999).

#### Second herbicide

To study the effect of binary herbicide mixtures and rotations, we consider two classes of herbicides used for Johnsongrass control, ACCase- and ALS-inhibitors (Peerzada et al. 2017). Due to the different sites of action, target-site resistance against herbicides of one class is uncorrelated with resistance against the other. However, in annual ryegrass (*Lolium rigidum*), multiple and cross-resistance to ACCase- and ALS-inhibitors has been reported (Torra et al. 2021) due to a combination of target-site and non-target-site resistance (Torra et al. 2021). Since we do not consider non-target-site resistance, we omit cross-resistance. The combined efficacy of the two herbicides applied as a mixture is calculated as *h_mix_* = 1*−*(1*−h*_1_)(1*−h*_2_) = *h*_1_ +*h*_2_ *−h*_1_*h*_2_, where *h*_1_ and *h*_2_ are the efficacies of the herbicides in mono-treatment (Bliss 1939). Likewise, suppose the effects of the fitness costs associated with the respective target-site resistance are independent. Then the fitness cost of a plant fully resistant against both herbicides is *c*_double resistant_ = 1*−*(1*−c*_1_)(1*−c*_2_) = *c*_1_+*c*_2_*−c*_1_*c*_2_. Here *c*_1_ and *c*_2_ are the fitness costs of resistance towards the individual herbicides.

### Stochastic simulations

Deterministic models capture selection as an evolutionary force, particularly for large populations. However, for small populations, strong selection may lead to the extinction of non-favoured types. Moreover, genetic drift acquires more relevance in small populations. We implemented stochastic simulations of our population-based model to account for such natural stochasticity. This approach allows for random shifts in allele frequencies and extinction. We provide the details of the algorithm in the Methods section, and the complete code is available on GitHub https://github.com/tecoevo/JohnsongrassDynamics for reproducibility.

## Results

### Genetic complexities

#### Standing genetic variation

We estimate the standing genetic variation for target-site resistance in untreated Johnsongrass by vector **v** in Eq. (22). The obtained population composition provided the initial seed bank and rhizomes composition for our deterministic population dynamics and corresponding stochastic simulations. We explore how the standing genetic variation for target-site resistance depends on the associated fitness cost (*c*) and its dominance (*k_c_*).

The frequency of resistance alleles R is controlled by the fitness cost on seed production (Fig. 2 a). It decreases with an increasing fitness cost. For fecundity reductions in resistant homozygotes exceeding 12 %, we expect a resistance allele frequency in the order of the considered de novo mutation rate *µ* = 10*^−^*^8^ (Haughn & Somerville 1987, Harms & DiMaio 1991); indicating that competition with the sensitive type prevents the establishment of the resistant mutants. A higher degree of dominance regarding the fitness cost decreases the R allele fraction slightly, which is less visible for small fitness costs. Likewise, the frequency of both resistant types is slightly reduced by a higher dominance of the fitness cost (Fig. 2 b). Remarkably, the expected frequency of heterozygotes shows little variation with the fitness cost. However, the expected frequency of resistant homozygotes increases with a decreasing fitness cost, most extreme for minimal fitness costs. This differential behaviour is because Johnsongrass is predominantly self-pollinated. Self-fertilised heterozygous plants (95 %) produce only one-half of heterozygous seeds and one-quarter of each homozygote. Therefore, we see an increased level of homozygosity under low resistance costs. For costs lower than 20 *−* 25 % (depending on the degree of dominance), we expect homozygous resistant plants to be more abundant than heterozygotes.

**Figure 2:**
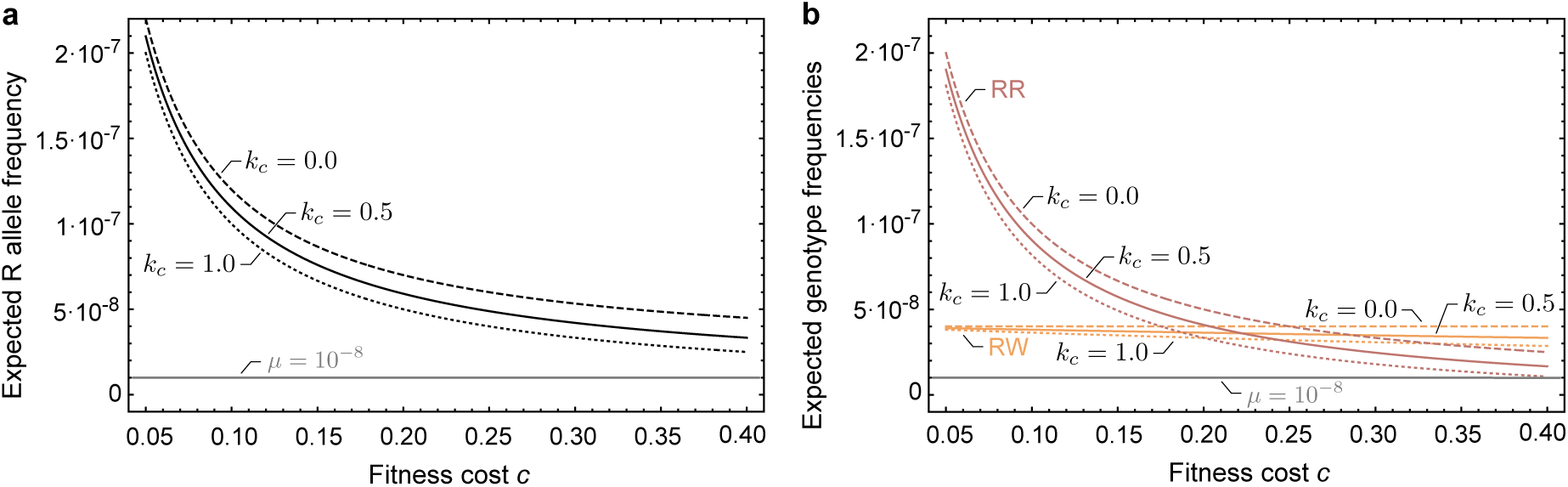
Variation of the approximated standing genetic variation for target-site resistance with the resistance cost. The genetic composition of untreated Johnsongrass populations is approximated on the basis of Eq. (22) and shown for different degrees of dominance (*k_c_*) of the resistance cost. Dashed lines correspond to a recessive resistance cost (*k_c_* = 0), solid lines indicate partial dominance (*k_c_*= 0.5) and dotted lines complete dominance(*k_c_* = 1). The grey line marks the spontaneous mutation rate of resistance alleles (*µ* = 10*^−^*^8^). **a**, Expected frequency of the resistance allele R in an untreated population depending on the resistance cost (*c*). **b**, Expected frequencies of the resistant genotypes in an untreated population as a function of the resistance cost (*c*). The frequency of resistant heterozygotes (RW) is shown in yellow and resistant homozygotes (RR) are represented in red.

In natural populations of annual ryegrass, plants with target-site resistance against an ALS-inhibitor were found at a frequency ranging from 1 *·* 10*^−^*^5^ to 1.2 *·* 10*^−^*^4^ (Preston & Powles 2002). Délye et al. (2013) detected one individual carrying a specific ACCase target-site resistance in a sample of 685 blackgrass (*Alopecurus myosuroides*) specimens collected prior to the introduction of ACCase-inhibiting herbicides. This abundance corresponds to a resistance allele frequency of 7.3 *·* 10*^−^*^4^. According to our model, we expect resistance alleles and resistant individuals in untreated fields at a frequency of 1 *·* 10*^−^*^5^ for a 0.1 % fitness cost on seed production (Data available on GitHub at https://github.com/tecoevo/JohnsongrassDynamics/tree/main/Fig2/data). Such a low fitness cost might not be detectable. Indeed, no associated fitness cost was detected for two out of three mutant ACCase alleles investigated by Menchari et al. (2008) in blackgrass. However, for a fitness cost on seed production of 30 %, reported for a mutant ACCase allele endowing herbicide resistance in Johnsongrass (Panozzo & Sattin 2021), the expected resistance allele frequency and cumulative frequency of resistant types range from 3.3 *·* 10*^−^*^8^ to 5.3 *·* 10*^−^*^8^ and 4.9 *·* 10*^−^*^8^ to 7.3*·*10*^−^*^8^, respectively, depending on the dominance of this fitness cost. This initial prevalence of target-site resistance is in the order of the spontaneous mutation rate (Haughn & Somerville 1987, Harms & DiMaio 1991) and considerably lower than what is reported in the literature (Preston & Powles 2002, Délye et al. 2013).

#### Resistance allele dominance and fitness cost dominance

An increase in the dominance of the resistance cost leads to a relatively small reduction in the overall fitness of heterozygotes, resulting in a minor decrease in the eventual abundance of heterozygotes and sensitive homozygotes (Fig. S1). Sensitive plants decrease in abundance with heterozygous plants due to the prevalence of self-pollination. Reduced resistance dominance drastically lowers heterozygotes’ fitness under herbicide application. The reduced survival strongly decreases the number of heterozygous and homozygous sensitive plants in the population.

A high dominance of resistance accelerates target-site resistance evolution when resistance is rare while hindering fixation of the resistance allele (Fig. S3 a). At low frequencies of target-site resistance, it is advantageous if heterozygotes have a high fitness and produce many offspring, of which most are resistant again. This, in turn, is a disadvantage for allele fixation since the sensitive allele is masked in heterozygotes.

#### Fitness cost

Using stochastic simulations, we investigate the impact of resistance cost on target-site resistance evolution and resulting population regrowth in herbicide-treated Johnsongrass. It is worth noting that the fitness cost determines the frequency of standing genetic variants, substantially affecting the probability of Johnsongrass escaping from control and regrowing.

Presumed that the resistant types manage to establish, the change in population composition is slightly faster under a low resistance cost on seed production (*c* = 0.001) compared to a high cost (*c* = 0.3) (Fig. 3 a). The resistant homozygotes outcompete other types more easily (see also Fig. S3 b). Moreover, with a low fitness cost of 0.1 %, the population escapes control by the herbicide in over half of the simulation runs (Fig. 3 b). Most of these simulated populations start to grow again within the first years (78.5 % within the first three years and 84.4 % within six years), where the year of escape from control and the start of regrowth is defined as the year in which the first homozygous resistant plants survive till reproduction. This resurgence is due to a high initial resistance allele frequency under low fitness costs (compare Fig. 2). Thus, resistant individuals will likely be in the field at the start of treatment. The impact of the fitness cost itself is comparably low. Under herbicide application, resistant plants have a major selective advantage over sensitive plants, even with a high fitness cost on seed production. For a higher fitness cost of 30 %, less than a quarter of simulated populations escape from control by the herbicide and regrow within 30 years (Fig. 3 b). The distribution of escapes from control is less skewed and more uniform, with almost no escapes in the first two years. Due to a low initial frequency of resistant types (compare Fig. 2), the population rescue depends on the probability of a new mutation arising or resistant seeds in the seed bank germinating.

**Figure 3:**
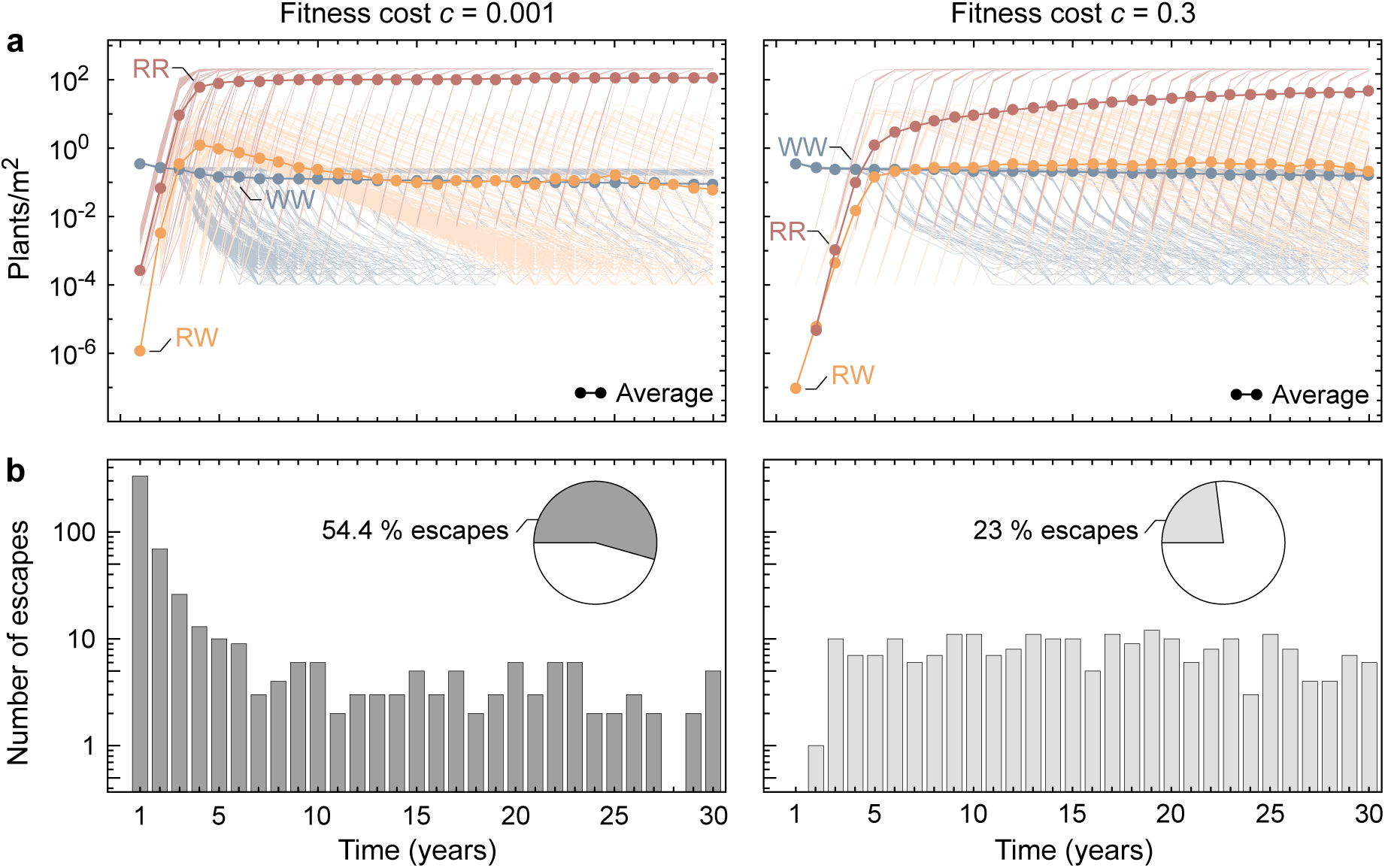
Simulated target-site resistance evolution and resulting population regrowth in herbicide-treated Johnsongrass for low and high resistance cost. Shown are the results of 1000 simulation runs obtained for a partially dominant resistance allele (*k_h_* = 0.5) and fitness cost (*k_c_*= 0.5). The initial genotype composition differs between the low (*c* = 0.001) and high (*c* = 0.3) fitness cost (compare Fig. 2 b). **a**, Changes in genotype composition of plants (*P*) over 30 years of herbicide application for low and high resistance cost. The frequency of sensitive plants (WW) is shown in blue, resistant heterozygotes (RW) in yellow and resistant homozygotes (RR) are represented in red. The thick lines with closed circles correspond to the average of all simulation runs, and the thin lines represent the individual realisations. **b**, Distribution of escapes from control over 30 years of herbicide application for low and high resistance cost. Weed populations can regrow under herbicide treatment if a resistant plant establishes on the field and reproduces. The year of escape from control is the year in which the first homozygous resistant plant survives till reproduction. The pie charts display the proportion of simulation runs where the weed population escapes from control and regrowths.

### Sexual reproduction

Genetic diversity mainly results from sexual reproduction, while perennials of a well-adapted type rapidly spread through asexual propagation. Two characteristics of Johnsongrass’ sexual reproduction, namely self-pollination and seed bank formation, seem particularly relevant for the population dynamics and target-site resistance evolution.

#### Self-pollination

Selfing increases the level of homozygosity in the population. Hence, the heterozygotes are significantly less abundant compared to a solely cross-pollinated population (Fig. S2), and the proportion of resistance alleles R increases faster under self-pollination (Fig. 4 b). The interplay of a higher generation of homozygous-resistant individuals and the selective pressure exerted by the herbicide causes the accelerated target-site resistance evolution in self-pollinated weed populations. It might seem surprising that our model predicts an initial decline in homozygous-resistant seeds under herbicide application if the weed population is cross-pollinating. This decline is because, in our model, the initial population composition derives under the assumption of no cross-pollination. The sensitive type dominates over the first years, producing a high fraction of sensitive pollen. Therefore, in a solely cross-pollinated population, pollination with sensitive pollen prevails.

**Figure 4:**
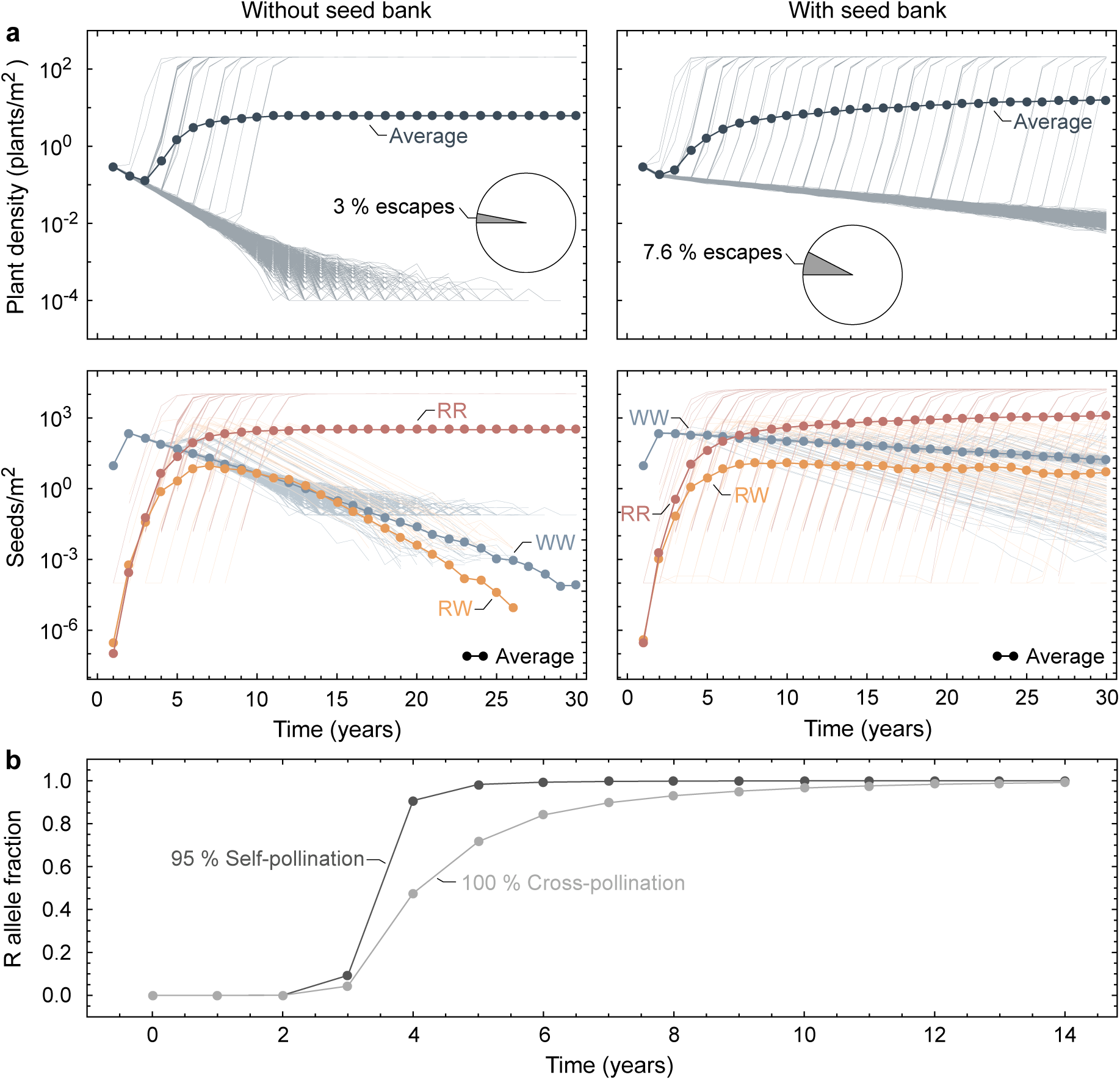
Predicted target-site resistance evolution in herbicide-treated Johnsongrass depending on seed bank formation and self-pollination. The results are obtained for a partially dominant resistance allele (*k_h_*= 0.5) and fitness cost (*k_c_* = 0.5). **a**, Simulated changes in Johnsongrass density (*P /A*) and genotype composition of seeds (in the seed bank (*B*) if formed, otherwise produced seeds (*S*) that survived the winter) over 30 years of herbicide application and tillage depending on the formation of a seed bank. Shown are the results of 1000 simulation runs. The thick lines with closed circles correspond to the average of all simulation runs, and the thin lines represent the individual realisations. The frequency of sensitive seeds (WW) is shown in blue, resistant heterozygotes (RW) in yellow and resistant homozygotes (RR) are represented in red. The pie charts display the proportion of simulation runs in which the weed population escapes from control and regrowths due to herbicide resistance evolution. **b**, Predicted changes in the frequency of the resistance allele R in Johnsongrass plants (*P*) under herbicide application for pure cross-pollination and 95 % self-pollination. Shown are predictions of our deterministic model.

#### Seed bank

Most simulated populations that lack a seed bank go extinct within 30 years under herbicide application and tillage (Fig. 4 a). At the same time, we see no extinctions within this period for simulated populations that include a seed bank. In the latter case, some resistant seeds might be left in the seed bank, even if the resistant types do not establish during the early years of herbicide application. The weed population thus has the potential to escape from control even after being controlled at very low densities for many years. Moreover, since populations with a seed bank are larger, the probability of mutations arising increases. Therefore, the distribution of escapes from control over time, observed in the stochastic simulations, is very wide for populations containing a seed bank. The escapes aggregate to 7.6 % of the realisations. In contrast, in simulations without a seed bank, populations started regrowing solely in the first nine years, accounting for 3 % of the runs.

Without a seed bank, the sensitive type disappears from almost all simulated populations, while most seed banks still contain some sensitive seeds after 30 years of recurrent herbicide application and tillage. The seed bank composition changes more slowly than the genotype composition of plants on the field, preserving genetic variation (see also Fig. S2). The seed bank only slightly delays the rise of resistance alleles in plants over several years of herbicide treatment (Fig. S3 c).

### Control strategies

#### Combined control regimes

We examined the impact of a binary herbicide mixture and the integration of soil tillage on population dynamics and target-site resistance evolution in Johnsongrass. The integration of tillage with herbicide application significantly reduces the weed population size (Fig. 5 a). This control improvement is caused by increased winter mortality of rhizomes and enhanced herbicide efficacy on rhizome shoots. Therefore, mutations are less likely to arise, and the proportion of simulation runs where the weed population regrows from resistant individuals also reduces. Nevertheless, the increased selective pressure with tillage causes the frequency of resistance alleles in the population to increase slightly faster than under herbicide application alone (Fig. 5 b). Our result contradicts the finding of Liu et al. (2019) that tillage delays the evolution of target-site resistance. However, we could qualitatively reproduce their result using a similar parameter set (see Fig. S4). Therefore, we acknowledge the deviation due to the significantly lower reproductive capacity implemented by Liu et al. (2019). The limited propagation increases the effect of higher rhizome mortality, overcoming the increased selective pressure on shoots under low frequencies of the resistance allele.

**Figure 5:**
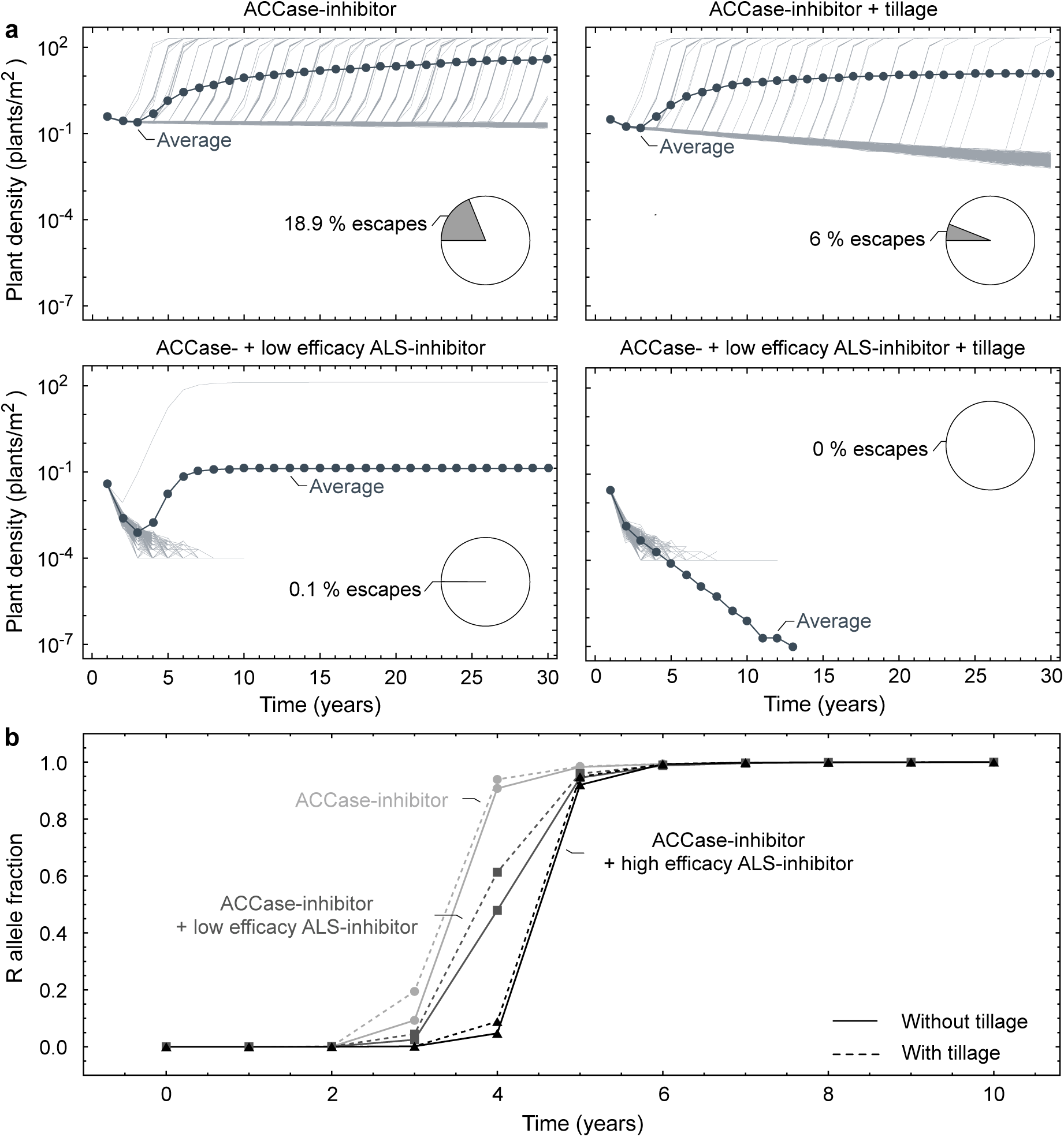
Predicted population dynamics and target-site resistance evolution in Johnsongrass under different control regimes. The results are obtained for a partially dominant resistance allele (*k_h_*= 0.5) and fitness cost (*k_c_* = 0.5). **a**, Simulated changes in Johnsongrass density (*P /A*) and proportion of populations escaping from control over 30 years of different control regimes. Shown are the results of 1000 simulation runs. The thick lines with closed circles correspond to the average of all simulation runs, and the thin lines represent the individual realisations. The pie charts display the proportion of simulation runs in which the weed population escapes from control and regrowths due to herbicide resistance evolution. The distinct control strategies are from left to right, top to bottom: ACCase-inhibitor application, ACCase-inhibitor application combined with tillage, application of ACCase-inhibitor and ALS-inhibitor with low efficacy, application of ACCase-inhibitor and ALS-inhibitor with low efficacy combined with tillage. **b**, Predicted changes in the frequency of the ACCase resistance allele R in Johnsongrass plants (*P*) under different control regimes. Shown are predictions of our deterministic model. The distinct control strategies are: ACCase-inhibitor (light grey line with closed circles) or ACCase-inhibitor and ALS-inhibitor with low (dark grey line with closed squares) or high (black line with closed triangles) efficacy applied solely (solid line) or combined with tillage (dashed line).

Adding a second herbicide with a reduced efficacy makes the simulated populations go extinct within 15 years and escapes very unlikely (Fig. 5 a). Furthermore, the second herbicide delays the resistance evolution against the first herbicide (Fig. 5 b). The reason is Johnsongrasses’ inherent ecology. As the herbicide efficacy is higher on seedlings than on rhizome shoots, the proportion of surviving plants originating from rhizomes increases when a second herbicide with a different mode of action is added. The selective pressure on these rhizome plants is lower than on plants that emerge from seeds. This effect is even more substantial for higher efficacies of the second herbicide. However, suppose a herbicide with reduced efficacy is applied individually. In that case, Johnsongrass’ exceptionally high reproductive capacity still allows the population to grow (Fig. S5). The high density, in turn, leads to de novo mutations conferring herbicide resistance.

#### Herbicide rotation

We further investigated the effect of the cycle length in binary herbicide rotations on the establishment of resistant plants on the field (Fig. 6). We consider rotations of two herbicides with different target sites and equal efficacies, where mixture and monotreatment can be viewed as the two extremes. Cycle length refers to the number of years one herbicide is applied recurrently before treatment switches to the other herbicide.

**Figure 6:**
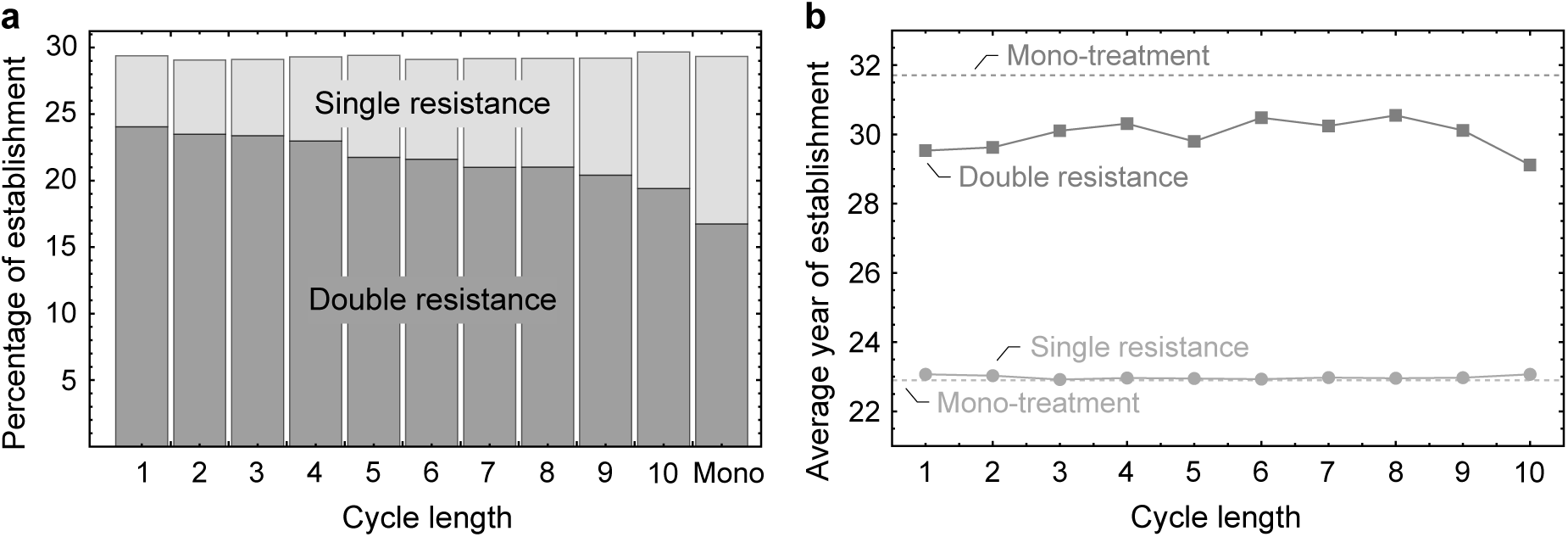
Simulated evolution of target-site resistance in Johnsongrass under binary herbicide rotations and mono-treatment. Shown are results of 10^5^ simulation runs obtained for a partially dominant resistance allele (*k_h_*= 0.5) and fitness cost (*k_c_* = 0.5). Considered are rotations of two herbicides with distinct target sites, ACCase- and ALS-inhibitors, Johnsongrass is known to develop target-site resistance towards. We assume equal efficacy here. The cycle length refers to the number of years one herbicide is recurrently applied before the treatment switches to the other herbicide. **a**, Proportion of simulated populations with resistant plants establishing on the field within 30 years of binary herbicide rotations depending on the cycle length and compared to mono-treatment. Bars show the percentage of simulation runs in which the plants (*P*) evolve resistance, with the light grey part referring to the population with single resistance, i.e. target-site resistance against only one of the herbicides, and the dark grey part reflecting double resistance. **b**, Average year of resistant plants establishing on the field within 30 years of binary herbicide rotations depending on the cycle length and compared to mono-treatment. The light grey line with closed circles displays the average year the first resistant plant survives till reproduction, and the dark grey line with closed squares depicts the average year of double resistant plants establishing on the field. The dashed lines show the reflective year of resistance occurrence under mono-treatment.

Herbicide rotations fail to reduce the probability of resistant plants establishing on the field. Varying slightly around 29.3 % observed in mono-treatment, the proportion of simulated populations with resistant plants is not dependent on the cycle length (Fig. 6 a). Due to the rapid population decrease, resistance evolves only in 0.02 % of the plants treated with a mixture of the two herbicides. In contrast, the emergence of double resistance decreases with increasing cycle length from 24 % in annual to 19.4 % in decennial rotation. In mono-treatment, we observe double resistant plants in 16.7 % of the simulated populations and none under the application of herbicide mixture. Resistance against one of the herbicides in a binary rotation is sufficient for the weed population to grow in the long run (compare Fig. S6). Further, herbicide rotations control sensitive plants with the same efficacy as a mono-treatment, leading to a similar probability of resistance adaptation. However, double resistance is costly for the plants, increasing the fitness cost on seed production in our model to 51 % compared to 30 % in single-resistant plants. When a herbicide is not applied, plants sensitive to this herbicide have a higher reproductive capacity. Therefore less frequent switching of herbicides reduces the risk of double resistance.

In our simulations, plants with single resistance established before the potential occurrence of double resistance. On average, the first resistant plants establish on the field after about 23 years of herbicide application, unaffected by cycling (Fig. 6 b). Under herbicide mixtures, double resistance occurs on average six to seven years afterwards, about two years earlier than under mono-treatment. Nevertheless, rotations of the herbicides can delay the resistance allele fixation due to the alternating selective pressure and the fitness cost associated with resistance (compare Fig. S7). Moreover, herbicide rotations delay weed population regrowth as compared to the application of a single herbicide (compare Fig. 5 b and Fig. S6). The average density reached by resistant populations after 50 years is 124 to 174 plants per m^2^ for herbicide rotations, considerably lower than 203 plants per m^2^ for mono-treatment.

## Discussion

Through developing a comprehensive life cycle model of Johnsongrass, we have addressed the goals set out in the Introduction: we showed, (i) that standing genetic variation for targetsite resistance is determined by the resistance cost, while its dominance has a minor impact; (ii) how high resistance dominance hastens the initial spread of resistance alleles but delays fixation due to the masking of sensitive alleles in heterozygotes; and the resistance cost affects probability and time of control escapes mainly by determining the standing genetic variation, with little effect of its dominance; (iii) that target-site resistance evolution is faster under self-pollination, and a seed bank can increase the probability of escape from control while maintaining genetic variation; (iv) that the integration of tillage with herbicide applications effectively reduces weed density and, thereby, the probability of escape from the control without delaying resistance evolution; how herbicide mixtures not only control Johnsongrass more effectively but also delay target-site resistance evolution due to the weeds inherent ecology; and that herbicide mixtures do not delay the onset of resistance, but can slow resistance evolution and delay weed population regrowth. Beyond the specific implications for Johnsongrass control, our results have significance in the broader context of the essential concepts in controlling weeds: life cycle details, chemical and physical control methods, resistance evolution and the prospects of ensuring food security. Below we draw out these conclusions.

Our analysis suggests that target-site resistance is associated with a meagre, probably undetectable, fitness cost when found in untreated populations at frequencies two orders of magnitude higher than the spontaneous mutation rate. For fitness costs exceeding 12 % on seed production, the resistance allele frequency in populations that never encountered the herbicide before is expected to be similar in magnitude to the mutation rate. Not all mutations endowing target-site resistance in weeds are known to involve a fitness cost (Menchari et al. 2008). Furthermore, target-site resistance against ALS- and ACCase-inhibitors were found with a natural frequency in the order of 10*^−^*^5^ to 10*^−^*^4^ (Preston & Powles 2002, Délye et al. 2013). According to our results, this would correspond to a meagre fitness cost. These findings coincide with the conclusions of Preston & Powles (2002). It is, therefore, likely that in Johnsongrass, targetsite mutations exist with very low or even no fitness cost. In this case, our simulations may underestimate the speed of resistance evolution and the risk of weed control failure.

We show a more than doubled probability of weed populations escaping herbicide control under a low fitness cost of 0.1 % compared to a fitness cost of 30 % on seed production. The latter cost was reported for a mutant ACCase allele endowing resistance in Johnsongrass (Panozzo & Sattin 2021). The expected change in population composition is faster under low resistance costs, and resistant plants establish earlier on the field, leading to population recovery. Thus, if the fitness cost of target-site resistance is low, control of the weed population with a single herbicide might fail quickly and should be avoided. Applying herbicide mixtures might be a good strategy in such a case, and integrating other control measures like tillage (Diggle et al. 2003, Peerzada et al. 2017).

Our results suggest that the fitness cost dominance has a minor impact on the prevalence of resistance and population dynamics under control. Unlike the fitness cost of resistance, herbicide resistance itself has no effect on the weed population in the absence of herbicides. The dominance of resistance, therefore, does not affect the standing genetic variation. However, it determines the fitness of heterozygotes under herbicide application, their abundance, and the sensitive type’s abundance. A high dominance of resistance initially increases the speed of resistance evolution. However, later the effect reverses due to the masking of sensitive alleles in heterozygotes, delaying resistance evolution. This effect is observed for diploids and tetraploid perennials (Holmes et al. 2022).

Especially, Holmes et al. (2022) found that target-site resistance alleles spread faster in haploids than in tetraploids since tetraploid plants need to acquire twice as many mutations to reach the same proportion of resistance alleles. However, in Johnsongrass, the W2027C target-site mutation was not found to be present on more than two ACCase alleles (Lorentz 2014). The cause might be a fitness penalty or the absence of homologous recombination between Johnsongrass’ two genomes (Lorentz 2014).

Our results indicate that target-site resistance evolution is faster in self-pollinating weed populations than in cross-pollinating populations. The increased homozygosity, combined with the herbicide’s selective pressure, causes a more rapid increase in the relative frequency of resistance alleles given resistant plants established. The higher number of homozygous-resistant individuals in the standing genetic variation of self-pollinated weeds comes at the cost of a lower number of resistance-carrying individuals, decreasing the probability of resistance adaptation for dominant target-site resistance. In another simulation study, populations of seed-propagated annual weeds needed to be larger for evolving resistance if they were self-pollinated compared to purely cross-pollinated populations (Diggle et al. 2003). However, the opposite is true for recessive target-site resistance. As heterozygotes are fully susceptible recessive target-site resistance is more likely to establish in a herbicide-treated field if the weed is self-pollinated. We derived our initial composition of seeds and rhizomes from Eq. (22) under the assumption of pure self-pollination. This genotype composition was used to calculate the population dynamics in an out-crossing population. However, we obtained similar results with a control-free initialisation phase of 30 years, allowing the population composition to stabilise under the assumed level of self-pollination.

Our simulations show that a seed bank significantly delays the extinction of a weed population under control. A seed bank preserves genetic variation and considerably increases the establishment probability of resistant mutants, leading to population regrowth. Therefore, we emphasise the necessity of considering the details of sexual reproduction and seed bank during risk assessment of control measures. It has been argued that herbicide resistance spreads mainly through asexual propagation (Liu et al. 2019). Once resistant plants have established, more resistant offspring might be generated via asexual propagation than seed production, considering the resistance cost on seed production and the possibility that produced seeds are not resistant. Resistance, however, generally originates in the seeds since mutations are most likely to arise during sexual reproduction. Even if standing genetic variants are present in the population, the number of seeds is considerably higher than that of rhizomes. Therefore, resistant individuals are more likely to be in the seed bank. Moreover, sexual reproduction is needed to generate homozygous-resistant offspring from heterozygous plants. We conclude that controlling the seed bank is essential in managing seed- and rhizome-propagated perennial weeds like Johnsongrass.

Integrating soil tillage with herbicide application further reduces Johnsongrass populations. We see no delay in the expected evolution of target-site resistance, as observed by Liu et al. (2019). Nevertheless, the probability of Johnsongrass regrowth is more than halved by the reduction in population size. This result agrees with field studies of Johnsongrass (McWhorter & Hartwig 1965) and blackgrass (Zeller et al. 2021). However, tillage increases the risk of soil erosion (Skøien et al. 2012) and diminishes soil water (Bescansa et al. 2006). Our study does not consider the risk of spreading resistant rhizomes and seeds with contaminated machinery.

Our results show herbicide mixtures control Johnsongrass most effectively, with minimal risk of weed populations escaping control. Including a second herbicide delays resistance evolution, with an increased effect for higher herbicide efficacies. These results agree with other simulation studies, demonstrating that herbicide mixtures effectively control resistance evolution and are superior to rotations (Diggle et al. 2003, Busi et al. 2020). These conclusions were derived assuming independent action of the herbicides, no cross-resistance and full application rates. Applying two herbicides at full rates includes a higher economic cost than a mono-treatment and increases the environmental impact. Also, possible mechanisms of cross-resistance and synergistic or antagonistic interactions of the respective herbicides need to be explicitly considered. A comprehensive understanding of the effect of herbicide mixtures could be attained by incorporating herbicide interactions and cross-resistance into the present model and investigating different application rates and ratios.

In our simulations, binary herbicide rotations failed to reduce the risk of target-site resistance compared to mono-treatments. Frequent herbicide switching even favours the evolution of double resistance. Due to the tremendous reproductive capacity of Johnsongrass, subpopulations resistant to only one of the herbicides will, in our model, still grow over a rotation period with equal application of both herbicides. Though not delaying the onset of resistance, herbicide mixtures can slow resistance adaptation. This aspect, however, only takes effect if a relevant cost of resistance induces a selective disadvantage of the resistant types in seasons the respective herbicide is not applied. Our findings agree with a simulation study of Busi et al. (2020), which found that binary rotations only delay resistance evolution if a fitness cost is assumed. However, they found more complex rotations, including four herbicides, to be more effective. Also, target-site resistance is not necessarily associated with a considerable fitness cost (Menchari et al. 2008). Moreover, rotations of two herbicides delay weed population regrowth. A field experiment on blackgrass found an average reduction in density of 59 % after seven years for yearly rotation of herbicide site of action compared to mono-treatment (Zeller et al. 2021). Our results show agreement with a simulation study by Diggle et al. (2003), in which rotations of herbicides were by far inferior to mixtures in controlling regrowth and herbicide resistance evolution. Overall our results promise an advantage of herbicide rotations over mono-treatment in terms of slowing population regrowth and target-site resistance evolution, presuming a fitness penalty, with no consistent advantage of fast or slow rotation. However, rotations do not reduce the resistance risk. Fast rotations even favour the occurrence of double resistance. In general, combinations of herbicides are far more effective in reducing herbicide resistance evolution and control failure.

Crop rotations, especially when they incorporate highly competitive crops like maize, can further reduce the population size of Johnsongrass (Scopel et al. 1988, Weisberger et al. 2019, Zeller et al. 2021). Holmes et al. (2022) studied the effect of crop rotations along with rotations of herbicides in a theoretical attempt. They highlight the effectiveness of a combination of crop rotation and rotation of herbicide classes in controlling Johnsongrass and delaying the evolution of herbicide resistance. However, they included resistance against only one of the herbicides, resulting from gene flow between resistant cultivated *Sorghum* and its wild relative. Our model can be extended to study the effect of crop rotations in terms of competition with the crop (Bargués-Ribera & Gokhale 2020). This extension can be achieved by adding the corresponding competition terms in the equations for density-dependant mortality (Eq. (9)) and reproduction (Eqs. (11) and (12)).

Besides standing genetic variation and de novo mutations, resistance can also arise from pollen-mediated gene flow from neighbouring resistant populations or closely related resistant crops and the introduction of herbicide-resistant rhizomes and seeds via contaminated equipment (Vrbničanin et al. 2017, Beckie et al. 2019). Maxwell et al. (1990) included seed immigration and gene flow from pollen in their population model of annual weeds, while Holmes et al. (2022) modelled resistance evolution in Johnsongrass with gene flow from resistant *Sorghum*. Extending our model to a landscape scale with intra- and inter-specific gene flow will comprehensively assess resistance evolution in herbicide-treated perennial weeds.

Overall, we have presented a theoretical framework capable of capturing the complex life cycle of perennial plants. The framework can be modified to model specific cases, as we have shown for Johnsongrass, and extended to answer pertinent questions from the points of view of different stakeholders in sustainable agriculture. For example, weed control measures always come with economic and socioecological facets. Thus, combining weed control and the associated socioeconomics in a single theoretical framework would be a fully functional tool built on our current model’s foundation.

## Methods

Routed in the biology of Johnsongrass or model describes the dynamics of a weed population growing in one field. We capture the perennial life cycle by considering five life-history stages: seeds, seedlings, rhizomes, shoots and adult plants. One time step of the model corresponds to one year. We implement stochastic simulations of this population-based model capturing the stochastic nature of mutations, genetic drift and extinctions. In this section, we describe the equations reflecting the plant reproduction and management interventions, as well as the stochastic simulations. We provide the derivation of the model parameters in the Supplementary Information with Table 1 showing all parameter values.

### Life cycle

Johnsongrass plants are dormant throughout the winter, overwintering as seeds or rhizomes in the ground (Warwick & Black 1983). At the beginning of a growing season, seeds germinate to produce seedlings, and shoots emerge from nodes on the rhizomes (Peerzada et al. 2017). Mature plants produce new rhizomes and, after flowering, viable seeds (Warwick & Black 1983, Peerzada et al. 2017). Figure 1 illustrates the life cycle of Johnsongrass and its implementation in the model with both reproduction pathways, sexual via seeds and asexual through rhizomes.

Initially, we neglect herbicide resistance and its genetics and concentrate solely on the plant life cycle. Rhizomes in the ground at the beginning of season *t* are denoted by *Z*(*t*). Not all axillary buds formed on the nodes of a rhizome develop as shoots; some develop into secondary rhizomes or stay dormant (Beasley 1970, Paterson et al. 2020). Let *b* be the number of nodes on a rhizome and *g_Z_* the proportion of nodes producing shoots during a season. Then the number of shoots (*T* (*t*)) at time *t* is given by,

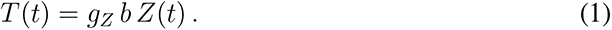

To begin with, we assume a constant seed bank *B* present in each season. Within a season *t*, only a certain proportion *g* of the seeds in the seed bank germinates to produce seedlings (*L*(*t*)) (Legutzamón 1986),

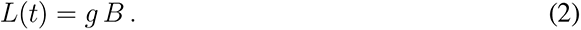

Seedlings and shoots grow up to adult plants (*P*). We do not distinguish adult plants by their origin from seeds or rhizomes as they show similar growth characteristics (McWhorter 1961, Horowitz 1972). Therefore,

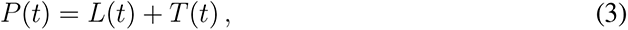

gives the number of adult plants (*P* (*t*)) present in season *t*. Plants form new rhizomes of which only a certain amount (1 *− d_Z_*) survives the winter to become the rhizomes of the next season *t* + 1,

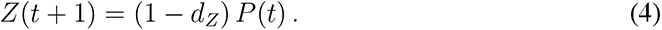

### Control and resistance

Our model includes herbicide application and soil tillage as weed management techniques. We consider target-site resistance endowed by a single resistance allele R in a diploid genome. We include spontaneous mutations to the resistance allele R in sexual reproduction but no back mutations. We assume the homozygous resistant genotype confers complete resistance against the herbicide. The factor 0 *≤ k_h_ ≤* 1 captures the dominance of resistance such that the herbicide controls WW plants with efficacy *h* and RW plants with efficacy (1 *− k_h_*) *h*. Due to incomplete control of the rhizomes by herbicides and resprouting from dormant rhizome buds, the herbicide efficacy on rhizome shoots *h_T_* is reduced compared to the herbicide efficacy on seedlings *h_L_* (Vidrine 1989). Soil tillage controls Johnsongrass rhizomes by exposing the fragments to low temperatures during winter, increasing mortality *d^∗^* (Warwick & Black 1983), and enhances herbicide control of rhizome shoots *h^∗^* (Hull 1970, Tuesca et al. 1999).

Let **T**(*t*) = (*T*_WW_(*t*)*, T*_RW_(*t*)*, T*_RR_(*t*)), **L**(*t*) = (*L*_WW_(*t*)*, L*_RW_(*t*)*, L*_RR_(*t*)), **P**(*t*) = (*P*_WW_(*t*)*, P*_RW_(*t*)*, P*_RR_(*t*)) and **Z**(*t*) = (*Z*_WW_(*t*)*, Z*_RW_(*t*)*, Z*_RR_(*t*)) be the vectors of shoot, seedling, plant and rhizome numbers in growing season *t*, respectively, and **B** = (*B*_WW_*, B*_RW_*, B*_RR_) the vector describing the composition of the seed bank. The population dynamics under herbicide application are then described by,

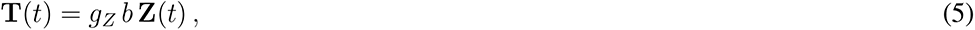

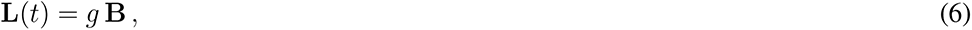

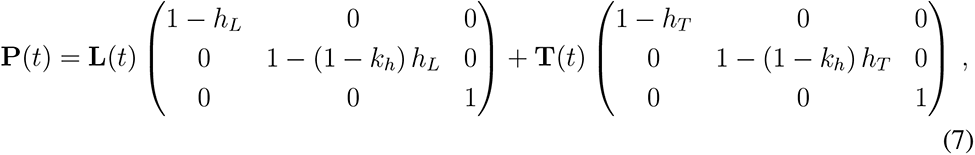

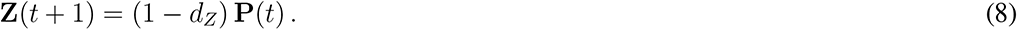

The herbicide efficiencies *h_L_* and *h_T_* are zero in a herbicide-free scenario. Enhanced winter mortality of rhizomes *d_Z_^∗^* depicts the use of mechanical control. For the case of the combined treatment, the herbicide efficacy on rhizome shoots *h_T_^∗^* is additionally increased.

### Intraspecific competition

Intraspecific competition affects the survival and the formation of reproductive structures in Johnsongrass (Williams & Ingber 1977, Schwinning et al. 2017). We describe the density-dependent mortality of Johnsongrass, resulting from intraspecific competition, by the hyperbolic self-thinning function

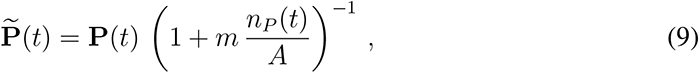

where **P**(*t*) is the vector of plants surviving till the end of season *t*, *A* the field size, *n* (*t*) =Σ*_i∈{_*_WW, RW, RR_*_}_ P_i_*(*t*) the total number of plants in the field without self-thinning (might be interpreted as young plants about to experience competition) and *m^−^*^1^ gives the highest possible density of Johnsongrass after self-thinning (Yoda et al. 1963). Therefore,

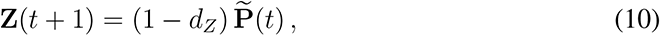

gives now the vector of rhizomes present at the start of season *t* + 1.

Comprehensive competition experiments in Johnsongrass are missing. Therefore, we base the fertility functions on the assumption that the total seed and rhizome yield per square meter approaches a constant value at moderate to high densities (Firbank & Watkinson 1985). Then the mean seed production (*f* (*t*)) and mean rhizome bud production (*b*(*t*)) per plant in season *t* can be described by

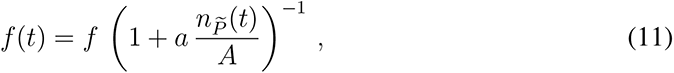

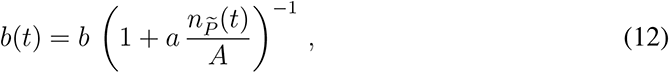

respectively, where *f* and *b* are the mean yields of an isolated plant, *a* is the area required by a plant to produce *f* seeds and *b* rhizome buds and *n_P_*_-_(*t*) =Σ*_i∈{_*_WW, RW, RR_*_}_ P_i_*(*t*) gives the total number of plants at the end of season *t* (after self-thinning) (Watkinson 1980). As the number of buds produced on a rhizome determines the number of shoots that can emerge from this rhizome in the next growing season,

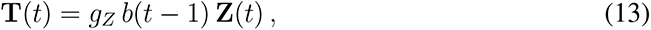

gives the shoots emerging in season *t*.

### Pollination and seed bank dynamics

We consider a fitness cost *c* on seed production associated with herbicide resistance (Panozzo & Sattin 2021). A dominance factor 0 *≤ k_c_ ≤* 1 controls the dominance of this fitness cost, such that the fecundity of RR and RS plants is reduced by *c* and *k_c_c*, respectively, compared to the fecundity *f* of susceptible plants. Johnsongrass is primarily self-pollinated (Warwick & Black 1983). Let *p_self_*denote the proportion of self-pollination. Then the number of seeds with genotype *i* (*S_i_^self^* (*t*)) produced during season *t* by self-pollinated plants can be calculated as,

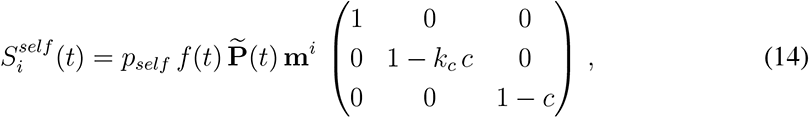

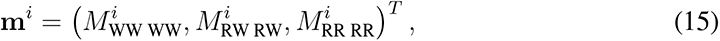

where *M_jk_^i^* gives the proportion of type *i* seeds produced by a plant of genotype *j* pollinated by type *k* pollen. The seed production follows Mendelian inheritance and includes spontaneous mutations to the resistance allele R occurring with probability *µ* (see Supplementary Information). However, up to 5 % of cross-pollination has been observed in fields with Johnsongrass plants growing at sufficiently small distances (Warwick & Black 1983). Over the *t*^th^ season a number of

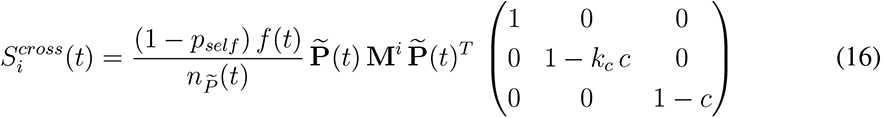

type *i* seeds (*S_i_^cross^*(*t*)) are produced by cross-pollinated plants, where

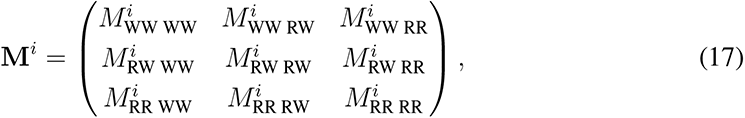

contains the proportions of type *i* seeds produced by the different matings, *M_jk_^i^*, *j, k ∈* {WW, RW, RR}. Adding up the seeds produced by self- and cross-pollinated plants gives the total number of genotype *i* seeds (*S_i_*(*t*)) produced in season *t*,

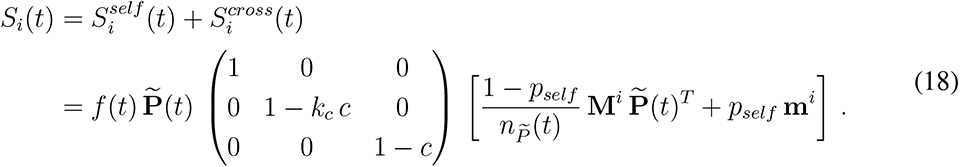

Johnsongrass seeds are dormant and form a seed bank in the ground (Warwick & Black 1983, Peerzada et al. 2017). Before new seeds enter the seed bank *B*, they are subject to post-dispersal predation and might lose viability or decay till the beginning of the next season with probability *d_S_*(Bagavathiannan & Norsworthy 2013). Within a season *t*, only a certain proportion *g* of seeds in the seed bank (*B*(*t*)) germinates to produce seedlings (*L*(*t*)),

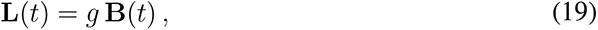

while non-germinated seeds may either be lost, due to decay and viability loss, with a probability *d_B_*or survive to be part of next season’s seed bank (Bagavathiannan & Norsworthy 2013). Therefore, the seed bank of season *t* + 1 consists of non-germinated seeds from the previous seed bank and the seeds produced during season *t*,

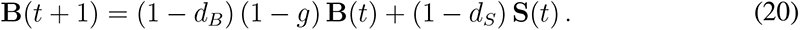

### Standing genetic variation

The natural frequency of target-site resistance exceeds the rate at which these mutations spontaneously occur, suggesting that herbicide resistance adaptation primarily proceeds from standing genetic variants (Kreiner et al. 2017). We derive a matrix model approximation of the entire life cycle dynamics. We use this matrix model to calculate the expected standing genetic variation for target-site resistance depending on the fitness cost associated with resistance and its dominance.

Let **P**(*t*) = (*P*_WW_(*t*)*, P*_RW_(*t*)*, P*_RR_(*t*)) and **B**(*t*) = (*B*_WW_(*t*)*, B*_RW_(*t*)*, B*_RR_(*t*)) be the vectors of plant and seed numbers in season *t*, respectively, and let (**P**(*t*), **B**(*t*)) be the population vector containing plant and seed numbers of all genotypes. We assume an infinite density limit (no self-thinning) and maximum reproduction (*f* (*t*) = *f, b*(*t*) = *b, ∀t ≥* 0). Further, we neglect cross-pollination (*p_self_* = 1) here. Under these assumptions, the population dynamics of Johnsongrass can be approximated by

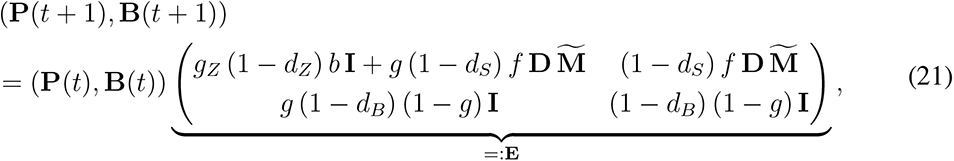

where **I** is the identity matrix of size 3,

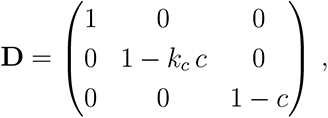

is the diagonal matrix incorporating the resistance cost on seed production and

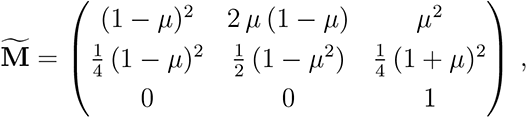

is the inheritance matrix, assuming simple Mendelian inheritance.

We use the *Perron-Frobenius* theorem 1 (see Supplementary Information) to derive the long-term population dynamics

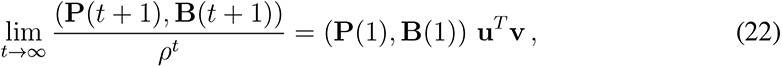

where *ρ* is the positive and simple eigenvalue of **E**, that is in absolute value the largest eigenvalue, and **u** and **v** are the corresponding positive right and left eigenvectors (Harris 1963). Thus, in the long run, the population composition is approximately a real multiply of **v**, regardless of the initial population. Therefore **v** gives us a rough estimate of the expected standing genetic variation for target-site resistance in a Johnsongrass population never exposed to the herbicide (see Fig. 2).

### Stochastic simulations

As in the underlying deterministic model, we model the population dynamics in discrete time steps of one growing season. The numbers of rhizome buds and seeds produced in season *t* by plants of a specific genotype *i* are drawn from Poisson distributions with mean *P_i_*(*t*) *b*(*t*) and *P_i_*(*t*) *f* (*t*), respectively. Probabilities, such as germination probability, death probabilities, and herbicide efficacy, are realised on the population level for the different genotypes using binomial distributed random numbers. The number of sensitive seeds germinating in season *t* is, for example, given by a binomial random number with parameters *B*_WW_(*t*) and *g*. We use multinomial random numbers to derive the initial seeds and rhizomes and to model fertilisation and mutation. Seeds resulting from self-pollination in heterozygous plants, for instance, are obtained from the multinomial distribution with the total number of seeds produced via self-pollination by heterozygous plants and the inheritance vector 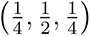 as parameters. For further details, we refer to the code available on GitHub.

## Data Availability

All data and plots are available on GitHub at https://github. com/tecoevo/JohnsongrassDynamics.

## Code Availability

All code is available on GitHub at https://github.com/tecoevo/ JohnsongrassDynamics.

## Acknowledgements

We thank Demetris Taliadoros for helpful discussions and acknowledge the generous funding from the Max Planck Society.

## Author Contributions Statement

Both authors conceived the study, developed the model, analysed the results and wrote the manuscript. D.L. formulated the mathematical model, the algorithm and ran the simulations.

## Supplementary Information

### Model parameters

#### Seed germination

Germination of Johnsongrass seeds is significantly influenced by temperature, light and burial depth (Horowitz 1972, Tóth & Lehoczky 2006, Krenchinski et al. 2015). Keeley & Thullen (1979) observed germination frequencies in the range of 24 to 91.5 % within three weeks for monthly sowing from March to October. In another study, no seed germination was documented at temperatures under 19*^◦^*C and a maximum of 64 % germination within 15 days was found at 39*^◦^*C (Horowitz 1972). However, Krenchinski et al. (2015) reported considerably lower germination frequencies, lower optimal germination temperature and reduced germination in the absence of light. They documented maximum germination at 30*^◦^*C of 18.50 and 13.75 % in the presence and absence of light, respectively. Seed germination of Johnsongrass decreases with increasing burial depth from reported 64 % germination from 1 cm depth to 6 % from a sowing depth of 25 cm (Tóth & Lehoczky 2006).

However, the population-level model does not account for these environmental factors. Warwick & Black (1983) found a mean seed germination frequency of 38 % in overwintering populations of Johnsongrass. This observation agrees with a study by Legutzamón (1986). Therein, 37 % of seeds left on the soil surface germinated within the first year and 25 % of seeds transferred from the surface to 15 cm depth in autumn, simulating tillage practice. Our germination probability *g* = 0.3 is between these values to account for seed burial due to natural causes or tillage. It is also consistent with the mean germination frequency documented by Egley & Chandler (1978).

#### Rhizome bud sprouting

In Johnsongrass, the development of rhizome buds is markedly controlled by apical dominance (Beasley 1970, Hull 1970). Rhizome apexes and emerged shoots partially suppress growth from adjacent buds (Beasley 1970, Hull 1970). Moreover, on branched rhizomes, secondary rhizomes influence the sprouting of axillary buds on the primary rhizome in an inverse manner (Beasley 1970). Bud sprouting and shoot growth in Johnsongrass are reported to be highly temperature dependant (Hull 1970, Horowitz 1972), with a threshold temperature of 10*^◦^*C, and to vary between populations (Loddo et al. 2012). In a study of Loddo et al. (2012) with 4-node rhizome segments, shoot emergence was highest at 16*^◦^*C, with a between population average of 48 %. Over the temperature range of 10 *−* 24*^◦^*C, a total of 26 and 39 % of the rhizome nodes developed sprouts in the two studied populations. Hull (1970) documented an average bud germination on one-node rhizome pieces of 14 and 92 % at 15 and 30*^◦^*C, respectively. However, Keeley & Thullen (1979) detected no relevant influence of temperature on rhizome bud sprouting (56 % at 18 *−* 35*^◦^*C) in their experiments, but a significant impact on shoot development. Shoots emerged from 13 and 88 % of the sprouted buds at 18 and 24 *−* 35*^◦^*C, respectively.

Our sprouting probability for buds on fragmented rhizomes in a tillage regime is 30 %, in agreement with the data of Loddo et al. (2012). Apical meristems, emerged shoots and lateral rhizomes can inhibit axillary bud development on branched rhizomes (Hull 1970). Therefore, we assume a reduced sprouting probability of 20 % for no-tillage regimes.

#### Maximum plant density

Johnsongrass produces tufts as a result of tillering, i.e. the development of secondary shoots from the crown area of existing plants (McWhorter 1971, Horowitz 1972). We do not explicitly model the production of tillers. However, the process of tillering is reflected in density-dependent reproduction. Lolas & Coble (1980) reported a considerable variation between different growing seasons in the number of tillers produced by a plant, ranging between 5 and 30 tillers per plant 60 days after planting, with an average of 19 tillers per plant grown from long rhizome pieces. Keeley & Thullen (1979) on average counted 5 to 20 tillers per plant, depending on the planting date. We assume that a plant not affected by intraspecific competition can, on average, produce 20 tillers.

In their experiments McWhorter (1971) observed an extreme variation between different Johnsongrass ecotypes in the number of culms growing in an area of one square meter, ranging from 65 up to 226 culms m*^−^*^2^. In another study, 200 Johnsongrass culms were, on average, counted per m^2^ in an untreated soybean field (Landry et al. 2016). Therefore we assume a possible highest density of 220 plants per m^2^ after self-thinning from very high densities. However, the ceiling density reached in populations growing from low densities is lower.

Tillers contribute to the total culm density on the field. However, high densities delay or even inhibit tillering in Johnsongrass (Williams & Ingber 1977). We assume that intraspecific competition becomes appreciable at a density of 10 plants per m^2^, i.e. 200 culms m*^−^*^2^ under maximum tillering. In the study of Keeley & Thullen (1979) Johnsongrass was planted at this density, meaning each plant had an area of 0.1 m^2^ to produce the reported yields.

#### Seed survival

Egley & Chandler (1978) documented no significant impact of the burial depth on seed longevity and an average of 62 % of seeds that were still viable after burial for 2.5 years. However, seeds were placed in polypropylene screen envelopes which prevented animal predation. It was not recorded if unrecovered seeds decayed or germinated. In another study, likewise excluding predation and not tracking germination, more prolonged seed survival in deep layers (15 and 22.5 cm) as compared to seeds in shallow layers (soil surface and 7.5 cm) was noticed (Legutzamón 1986). After three years, they recovered less than 15 % of viable seeds from the shallow layers, while 34.5 to 53.8 % of seeds in the deeper layers were still viable. Nevertheless, regardless of the burial depth, virtually no viable seeds were found in their experiments after six years. Moreover, Legutzamón (1986) carried out one experiment in bare soil with different treatments. Some treatments comprised regular changes between the soil surface and 15 cm depth to examine the effect of cultivation on seed longevity. They found maximum seed survival to be less than 2.5 years. However, they discussed that the absence of vegetation in that experiment might have contributed to a faster loss of viable seeds. Bagavathiannan & Norsworthy (2013) reported 64 to 75 % of seed loss due to predation, from dispersal in autumn till next spring, when seeds were left on the soil surface (no-tillage). Most recovered seeds (78 *−* 91 %) were buried in the soil. In the same period, they documented 6 to 17 % seed decay. Also, a loss in viability of 23 % was observed that increased to 52 % (including germination) till the subsequent autumn. Overall the mean proportion of active seed bank still present in spring, after seed loss due to predation, decay and loss in viability, was 10 to 11 % and reduced to 7 to 8 % in autumn, after subsequent seed loss through decay, loss in viability and germination (Bagavathiannan & Norsworthy 2013). They documented no significant impact of residue cover or burial depth on seed decay and viability loss.

We consider only post-dispersal seed predation, as most surviving seeds are then buried in the soil and therefore subsequently inaccessible for predators, at a proportion of 75 % (Bagavathiannan & Norsworthy 2013). Our model includes 17 % seed decay and 23 % loss in viability from the time of seed dispersal in autumn till next spring. We choose 24 % and 31 % yearly seed loss due to decay and loss in viability, respectively. With that, seed loss in our model is in the range reported by Bagavathiannan & Norsworthy (2013), but higher than found in the undisturbed experiments of Egley & Chandler (1978) and Legutzamón (1986). However, our parameters are in good agreement with the simulated cultivation experiments conducted by Legutzamón (1986). Bagavathiannan & Norsworthy (2013) calculated the proportion of predation, correcting for the proportion of decayed seeds. Following this, viable freshly produced seeds are lost to 0.94 % (1 *−* (1 *−* 0.75 *−* 0.17)(1 *−* 0.23) *≈* 0.94) in our model, and the yearly loss of viable seeds in the seed bank is 48 % (1 *−* (1 *−* 0.24)(1 *−* 0.31) *≈* 0.48).

#### Rhizome survival

Winter mortality of Johnsongrass rhizomes is highly dependent on the soil temperature, determined by climatic conditions and burial depth (Stoller 1977, Warwick et al. 1986, Hartzler et al. 1991). Stoller (1977) documented about 5 % of rhizomes surviving at a burial depth of 20 cm and no survival above 20 cm depth in the first winter with a temperature record of *−*9*^◦^*C at 20 cm. In the second milder winter, more than 60 % of the rhizomes survived at 20 cm and rhizome survival increased with increasing depth in soil up to over 90 %. For overwintering populations, Warwick et al. (1986) reported a dramatic reduction of rhizome survival at 20 cm depth to 20 % in a more northern climate compared to 97 % survival under milder climatic conditions. They also recorded a survival dependency on burial depth, with only one out of 16 rhizomes surviving at a burial depth of 10 cm and 87.5 % winter survival at 30 and 50 cm. Hartzler et al. (1991) also found rhizome survival directly related to burial depth. With 5 and 25 % survival of 7.5 and 15 cm long rhizome pieces buried in soil 6 cm deep, survival of longer rhizome pieces was higher. Simultaneously, they did not notice a significant difference at 25 cm depth, where survival ranged between 85 and 87 %. Most rhizome biomass and nodes are found in the upper 30 cm, with the highest amount between 7.5 and 15 cm (Hartzler et al. 1991).

Our model does not account for the depth of rhizome burial or climatic conditions. We use the rhizome survival and distribution data published by Hartzler et al. (1991) to calculate the average winter mortality of long rhizome segments *d_Z_* = 0.35, used in our model as the winter mortality in no-tillage regimes. Tillage cuts Johnsongrass rhizomes into smaller pieces and partly transports them to the soil surface. We assume that under control regimes comprising tillage, the share of rhizome pieces in the shallow layer is increased by 10 %. Together with the rhizome fragmentation, this leads to higher average winter mortality of *d_Z_* = 0.6.

All non-germinated rhizomes rotted in the experiments of Warwick et al. (1986). Furthermore, after sprouting, the rhizomes of Johnsongrass disintegrate within 19 to 24 days (McWhorter 1961). Therefore we only consider winter survival and assume that rhizomes can only give rise to shoots in the subsequent growing season.

#### Herbicide efficacy

The herbicide efficacy on Johnsongrass aboveground material depends on the developmental stage of the plants, with small young plants killed with higher efficacy than larger older plants (Tuesca et al. 1999, Dogan & Boz 2005, Johnson & Norsworthy 2014). The efficacy also varies between years due to environmental factors and application timing (Vidrine 1989, Tuesca et al. 1999).

Johnson et al. (2003) and Johnson & Norsworthy (2014) report the investigated ACCase-inhibitors to have a higher efficacy on Johnsongrass than ALS-inhibitors included in their studies. A single application of the ACCase-inhibitor Quizalofop led to a 97 % reduction in Johnsgrass biomass three weeks after treatment. In contrast, single applications of the ALS-inhibitors Nicosulfuron and Primisulfuron reduced fresh weight biomass by only 88 % (Johnson et al. 2003). In the second study, a higher rate of the ACCase-inhibitor Clethodim controlled small plants (15 and 30 cm) at 97 % two weeks after treatment while Nicosulfuron provided 90 % control of 15 cm Johnsongrass and only 74 % control of 30 cm plants (Johnson & Norsworthy 2014). Even though they noticed Johnsongrass regrowth after two weeks, stand reduction four weeks after treatment, compared to an untreated control, was similar to the initial weed control.

Due to insufficient translocation of active ingredients and the inactive state of rhizome nodes, rhizomes are controlled inadequately (Beasley 1970). Regrowth from the latter may occur, resulting in a reduced herbicide efficacy on rhizome shoots (Beasley 1970, Tuesca et al. 1999). In a study of Vidrine (1989), the average control of Johnsongrass seedlings provided by different ACCase-inhibiting herbicides was 99 % three weeks after treatment and 95 % for rhizome shoots. Sethoxidim failed to control Johnsongrass shoots effectively in one year of the experiment. Excluding sethoxydim, the other three ACCase-inhibitors controlled seedlings and shoots at 99.8 and 98.5 %, respectively. Johnsongrass control was reduced 12 weeks after treatment. Nevertheless, with one exception, control was still at least 86 % and, in most cases, even above 90 %. Winton-Daniels et al. (1990) reported that regrowth occurred after all treatments with ACCase-inhibitors but that sequential treatments typically control Johnsongrass better than single applications. A second herbicide application later in the season reduces the time for Johnsongrass plants to regrow. Therefore, regrown plants might be unable to produce new rhizomes and viable seeds until the end of the growing season. Given the above points, it is reasonable to assume that the final control provided by a sequential treatment is similar to the initial control a single application gives. As per Vidrine (1989), we choose the efficacy of ACCase-inhibiting herbicides to be 99.8 % on seedlings and to be reduced on shoots with 98.5 %. In some parts of our manuscript, this efficacy is also assumed for the ALS-inhibitor to secure effective weed reduction, for example, in herbicide rotations, and is denoted by ’high efficacy’. As ALS-inhibiting herbicides are reported to control Johnsongrass with a lower efficacy (Johnson et al. 2003, Johnson & Norsworthy 2014), we also investigate a low ALS-inhibitor efficacy of 92 and 90 % on seedlings and rhizome shoots, respectively, denoted by ’low efficacy’. Mechanical practices inducing increased bud activity as a response to rhizome fragmentation allow for improved control of rhizomes and shoots by herbicide application (Hull 1970, Tuesca et al. 1999). Therefore herbicide efficacy on shoots of fragmented rhizomes is assumed to be intermediate between seedlings and shoots on intact rhizomes.

#### Inheritance

We suppose simple Mendelian inheritance. Mutations to the resistance allele R are assumed to arise with probability *µ* = 10*^−^*^8^, but back mutations are not considered (Haughn & Somerville 1987, Harms & DiMaio 1991). Therefore the inheritance matrices (17) used to calculate the number of seeds arising from cross-pollination (16) are given by,

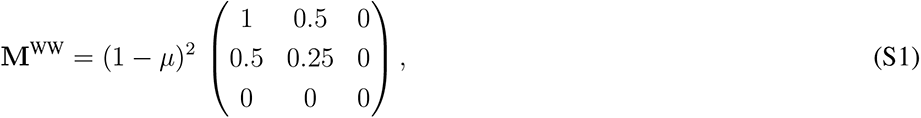

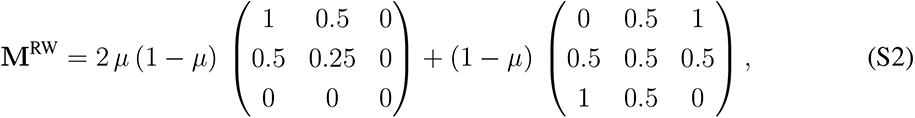

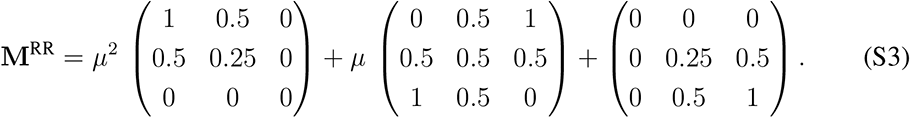

### Matrix theory

We make use of a well-known result from the theory of matrices.

#### Theorem 1 (Perron-Frobenius (Harris 1963))

*Let* **A** *be a nonnegative k x k matrix, such that* **A**^N^ *is positive for some N ∈ N. Then* **A** *has a positive and simple eigenvalue ρ that is in absolute value greater than any other eigenvalue of* **A**. *ρ corresponds to positive right and left eigenvectors* **u** *and* **v**, *which are the only nonnegative eigenvectors of* **A**. *Moreover, for n ∈ N we have*

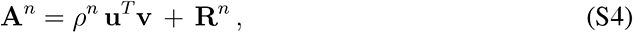

*with the normalisation* **u v**^T^ = 1. *Furthermore,* 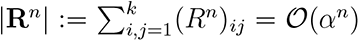 *for some α,* 0 *< α < ρ*.

From Eq. (A.4) it follows that in the limit

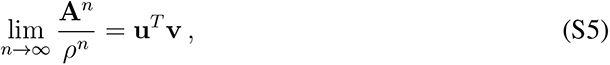

holds, i.e. **A***^n^* is dominated by *ρ^n^* for *n → ∞*.

---

**Fig. S1:**
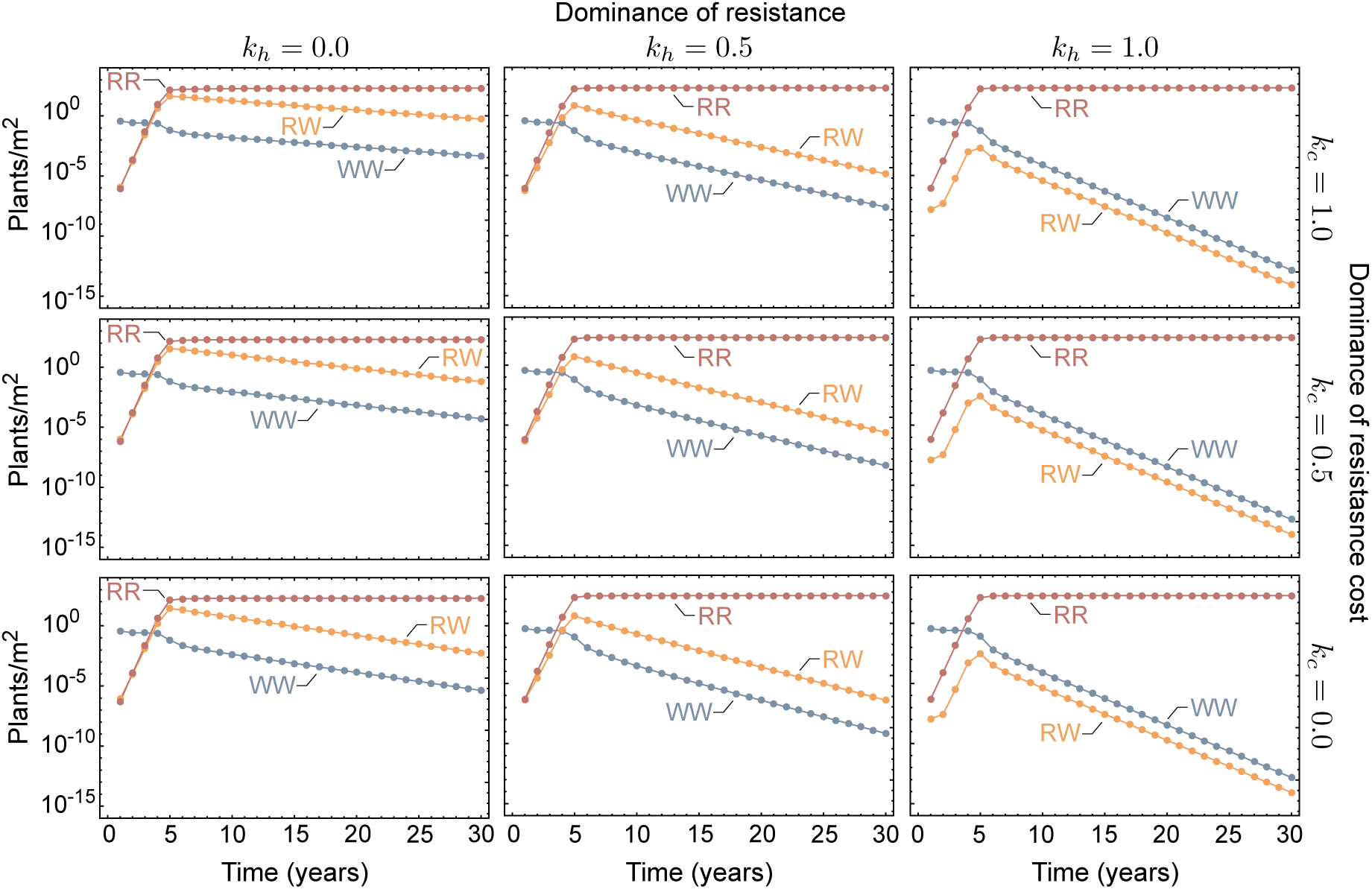
Predicted target-site resistance evolution in herbicide-treated Johnsongrass depending on the resistance and cost dominance. Shown are predictions of our deterministic model for the genotype composition of Johnsongrass plants (*P*) over 30 years of herbicide application for different degrees of dominance regarding resistance (**k_h_**) and the fitness cost associated with resistance (**k_c_**). (*k* = 0) corresponds to recessivity, (*k* = 0.5) indicates partial dominance and (*k* = 1) complete dominance. The frequency of sensitive plants (WW) is shown in blue, resistant heterozygotes (RW) in yellow and resistant homozygotes (RR) are represented in red. The initial genotype composition differs with the dominance of resistance and fitness cost.

**Table S1:**
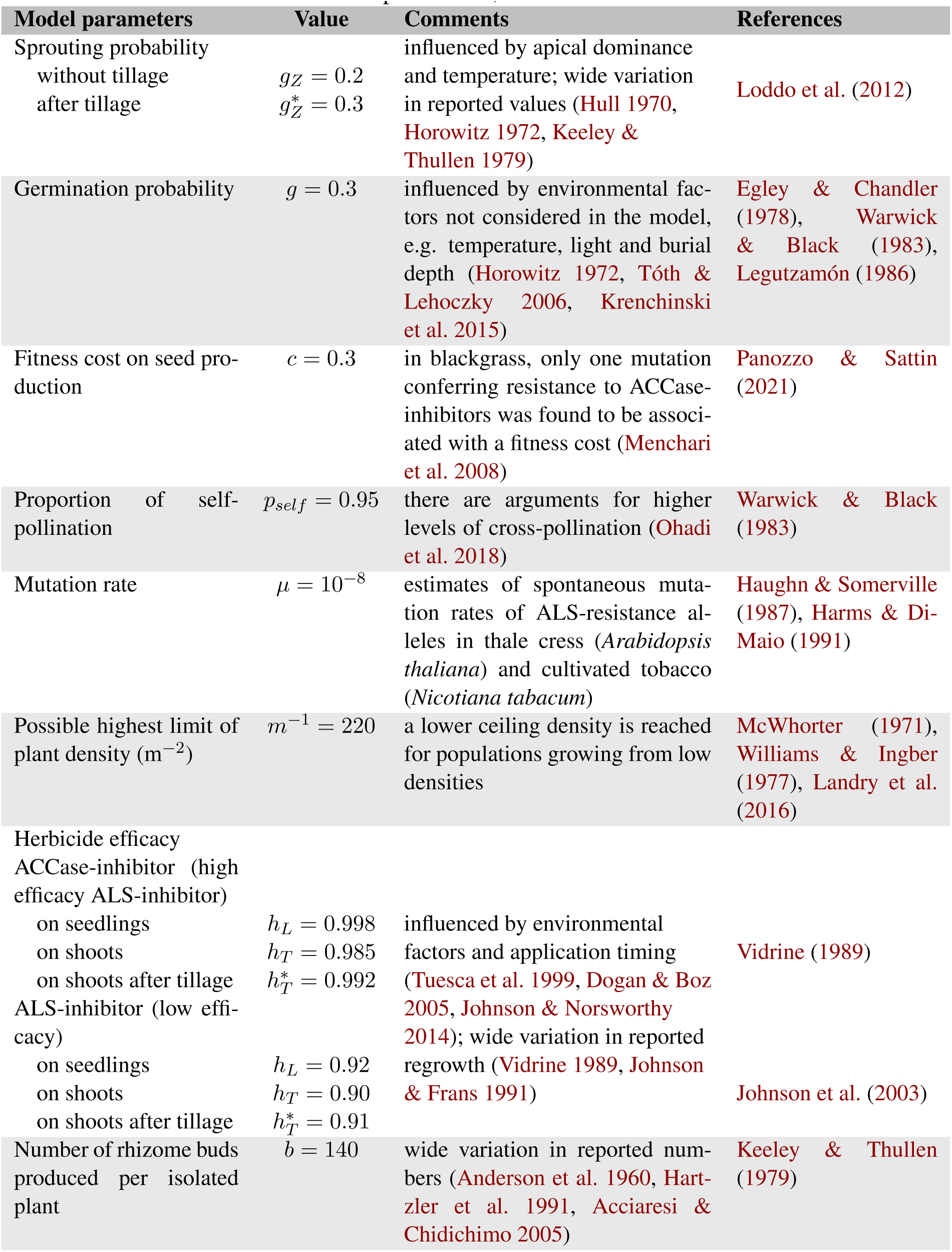
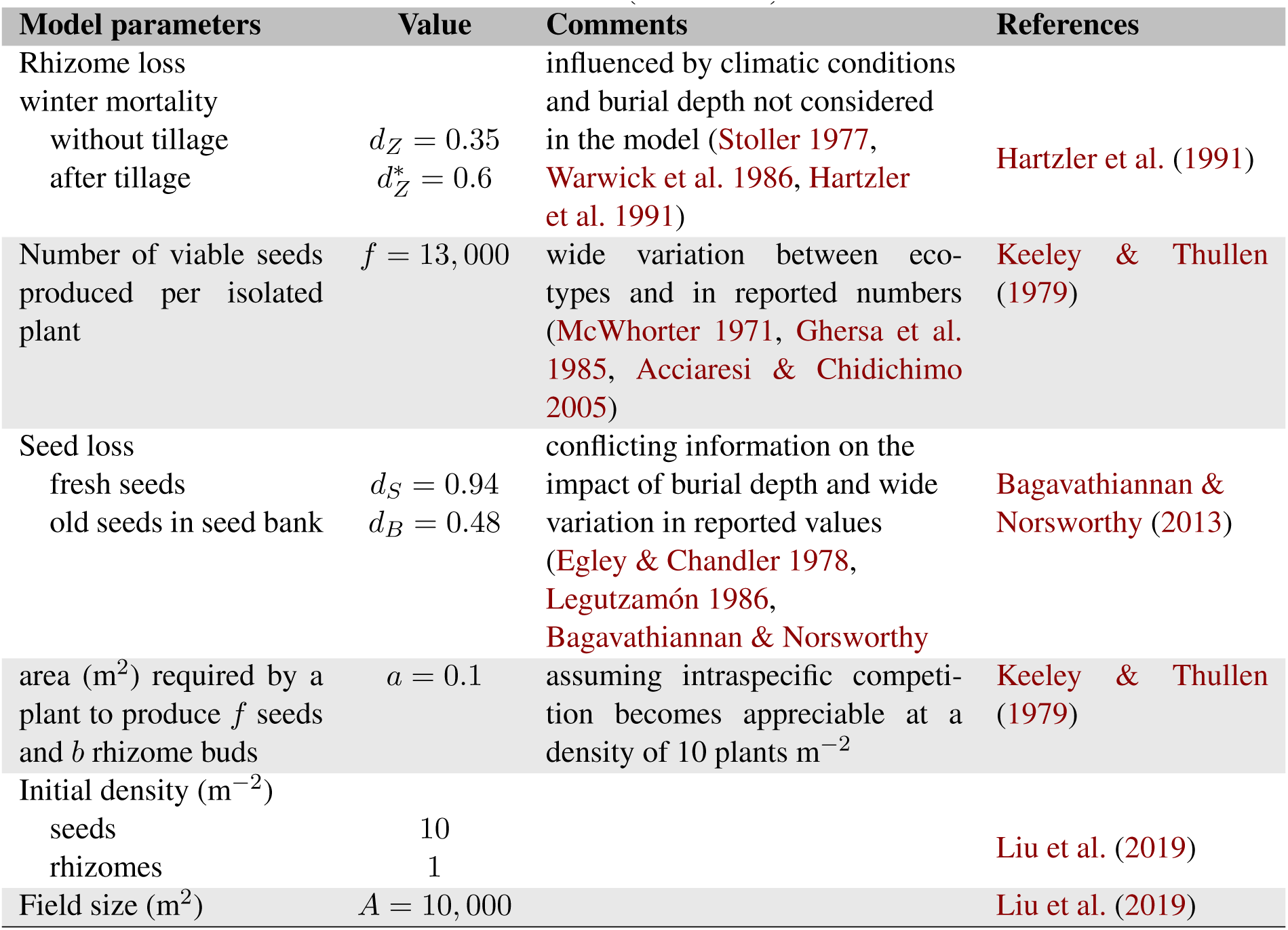
Model parameters, values and references.

**Fig. S2:**
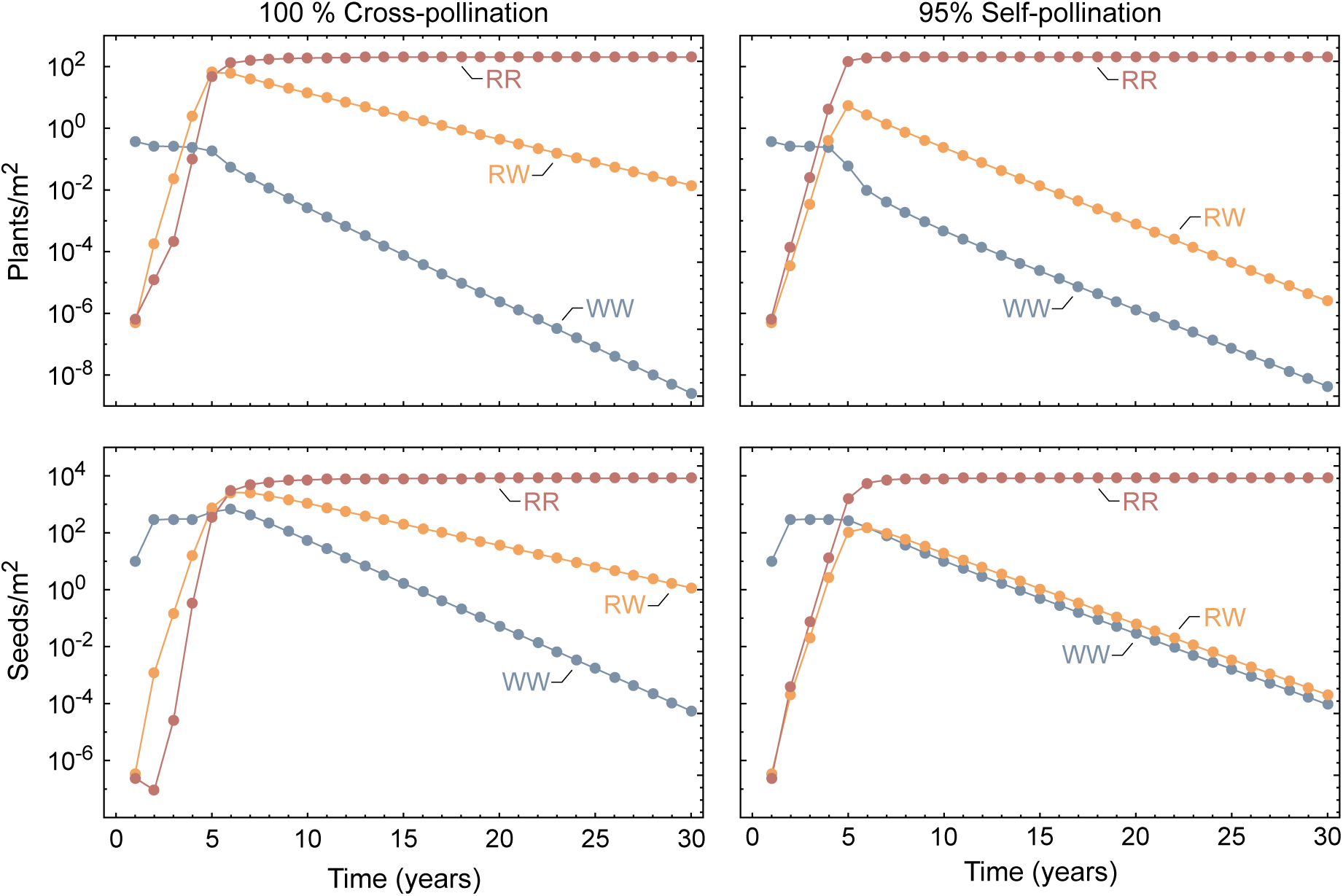
Predicted target-site resistance evolution in herbicide-treated Johnsongrass depending on self-pollination. The results are obtained for a partially dominant resistance allele (*k_h_* = 0.5) and fitness cost (*k_c_* = 0.5). Shown are predictions of our deterministic model for the genotype composition of Johnsongrass plants (*P*) and seeds (*B*) over 30 years of herbicide application for pure cross-pollination and 95 % self-pollination. The frequency of sensitive plants (WW) is shown in blue, resistant heterozygotes (RW) in yellow and resistant homozygotes (RR) are represented in red.

**Fig. S3:**
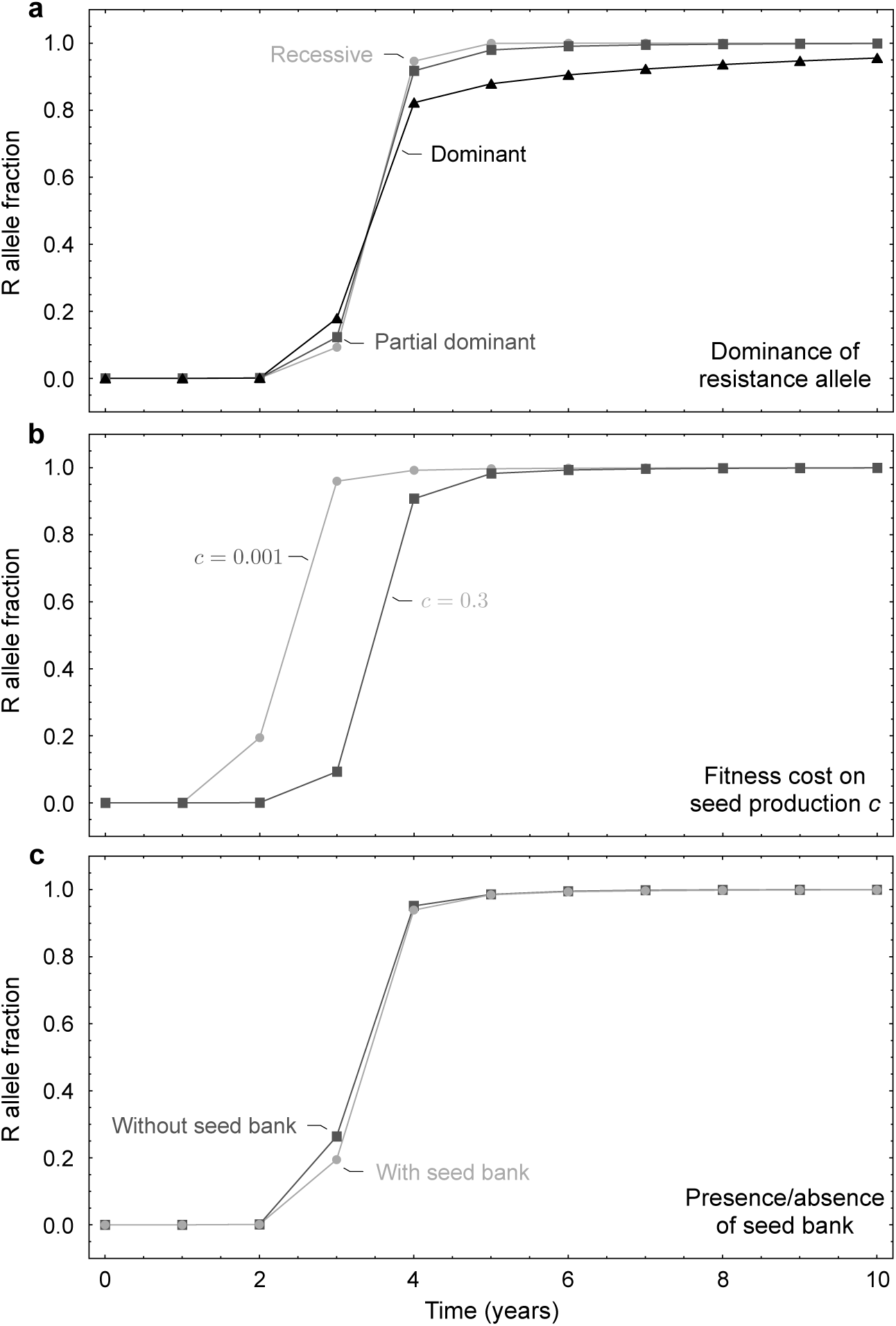
Predicted target-site resistance evolution in herbicide-treated Johnsongrass for different degrees of resistance dominance, low and high resistance cost and depending on seed bank formation. Shown are predictions of our deterministic model obtained for a partially dominant resistance allele (*k_h_*= 0.5) (varies in **a**) and fitness cost (*k_c_* = 0.5). **a**, Changes in frequency of the resistance allele R in Johnsongrass plants (*P*) under herbicide application for different degrees of resistance dominance (*k_h_*). Where recessive corresponds to *k_h_* = 0 (light grey line with closed circles), partially dominant to *k_h_*= 0.5 (dark grey line with closed squares) and dominant to *k_h_* = 1 (black line with closed triangles). **b**, Changes in frequency of the resistance allele R in Johnsongrass plants (*P*) under herbicide application for low (*c* = 0.001) and high (*c* = 0.3) resistance cost. The initial genotype composition differs between the low (light grey line with closed circles) and high (dark grey line with closed squares) fitness cost (compare Fig. 2 b). **c**, Changes in frequency of the resistance allele R in Johnsongrass plants (*P*) under herbicide application depending on the formation of a seed bank. The light grey line with closed circles c_4_o_5_rresponds to a population with a seed bank and the dark grey line with closed squares to a population without a seed bank.

**Fig. S4:**
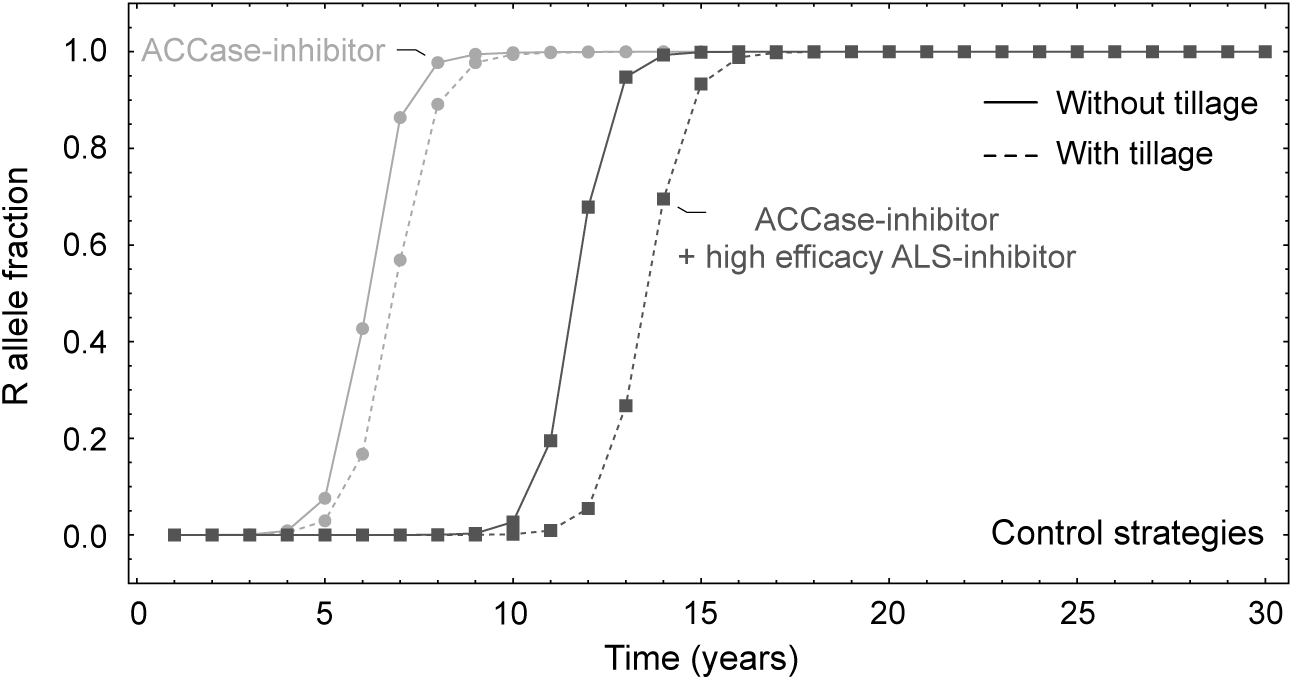
Predicted target-site resistance evolution in herbicide-treated Johnsongrass for different control regimes. Shown are deterministic predictions for the changes in the frequency of the resistance allele R in Johnsongrass plants (*P*) under different control strategies. The distinct control strategies are: ACCase-inhibitor (light grey line with closed circles) or ACCase-inhibitor and ALS-inhibitor with equal efficacy (dark grey line with closed squares) applied solely (solid line) or combined with tillage (dashed line). These dynamics where obtained with a different version of our population-based model. Compared to the model presented in this manuscript, this model does not include density dependence in reproduction and no mortality of seeds before entering the seed bank. Furthermore, the parameter set is based on the one used by Liu et al. (2019) along with a yearly seed bank mortality of 20 %. The implementation with all parameter values is available on GitHub at https://github.com/tecoevo/JohnsongrassDynamics.

**Fig. S5:**
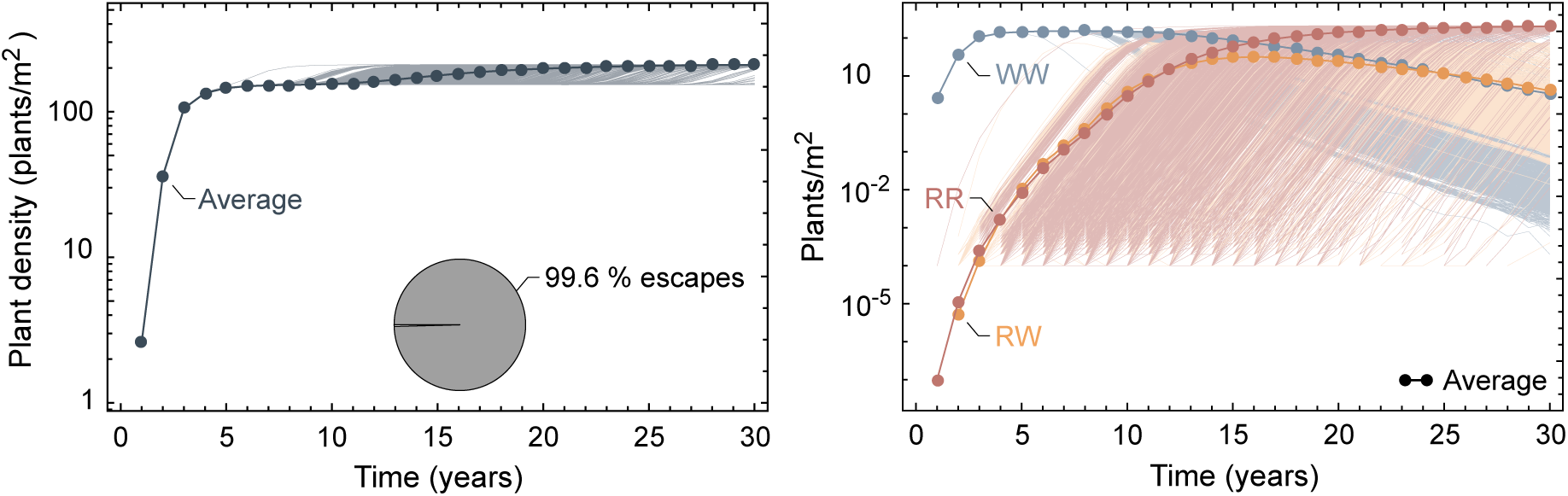
Simulated population dynamics and target-site resistance evolution in Johnsongrass treated with low efficacy ALS-inhibitor. Shown are changes in Johnsongrass density (*P /A*) and genotype composition of plants (*P*) over 30 years of ALS-inhibitor application with low efficacy of 1000 simulation runs obtained for a partially dominant resistance allele (*k_h_* = 0.5) and fitness cost (*k_c_*= 0.5). The thick lines with closed circles correspond to the average of all simulation runs, and the thin lines represent the individual realisations. The frequency of sensitive seeds (WW) is shown in blue, resistant heterozygotes (RW) in yellow and resistant homozygotes (RR) are represented in red. The pie charts display the proportion of simulation runs where the weed population escapes from control and regrowths due to herbicide resistance evolution.

**Fig. S6:**
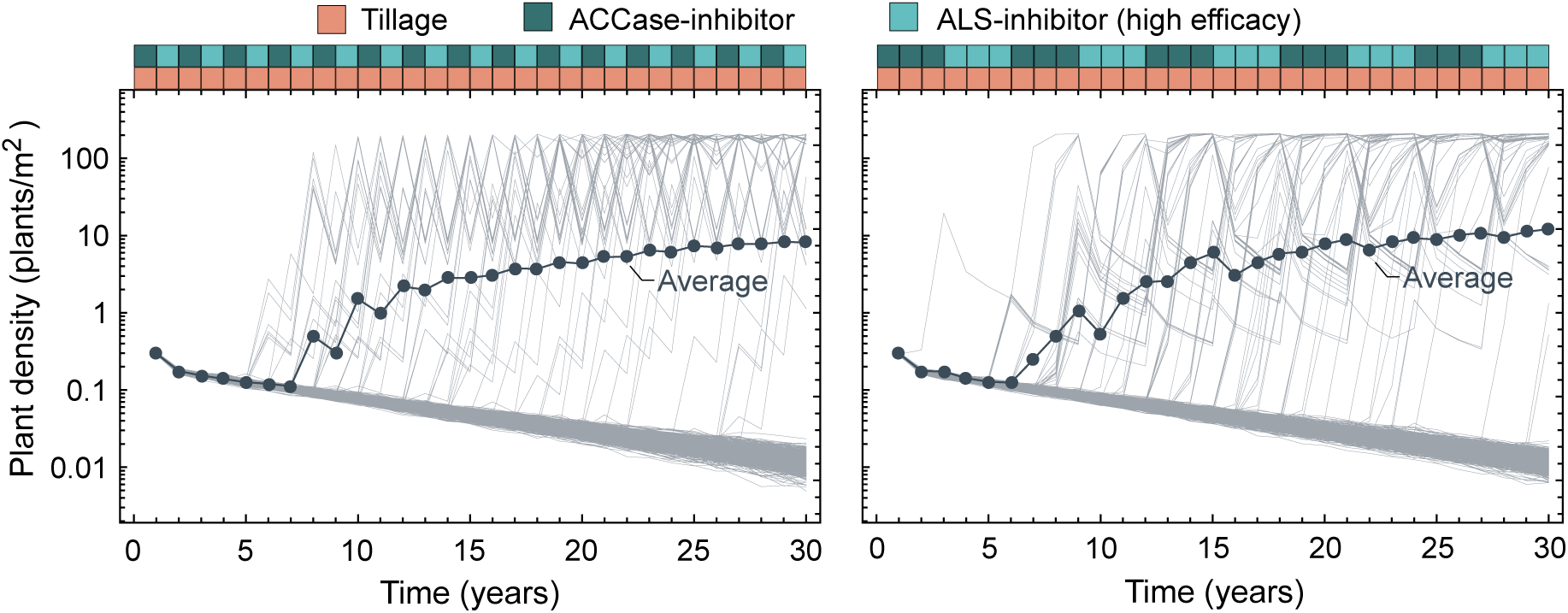
Simulated population dynamics in Johnsongrass for annual and three-annual herbicide rotations. The simulation results are obtained for a partially dominant resistance allele (*k_h_* = 0.5) and fitness cost (*k_c_*= 0.5). The left panel shows changes in Johnsongrass density (*P /A*) over 30 years for a yearly rotation of ACCase- and ALS-inhibitor combined with soil tillage. Population dynamics under a rotation of the same herbicides with a cycle length of three years combined with tillage are displayed in the right panel.

**Fig. S7:**
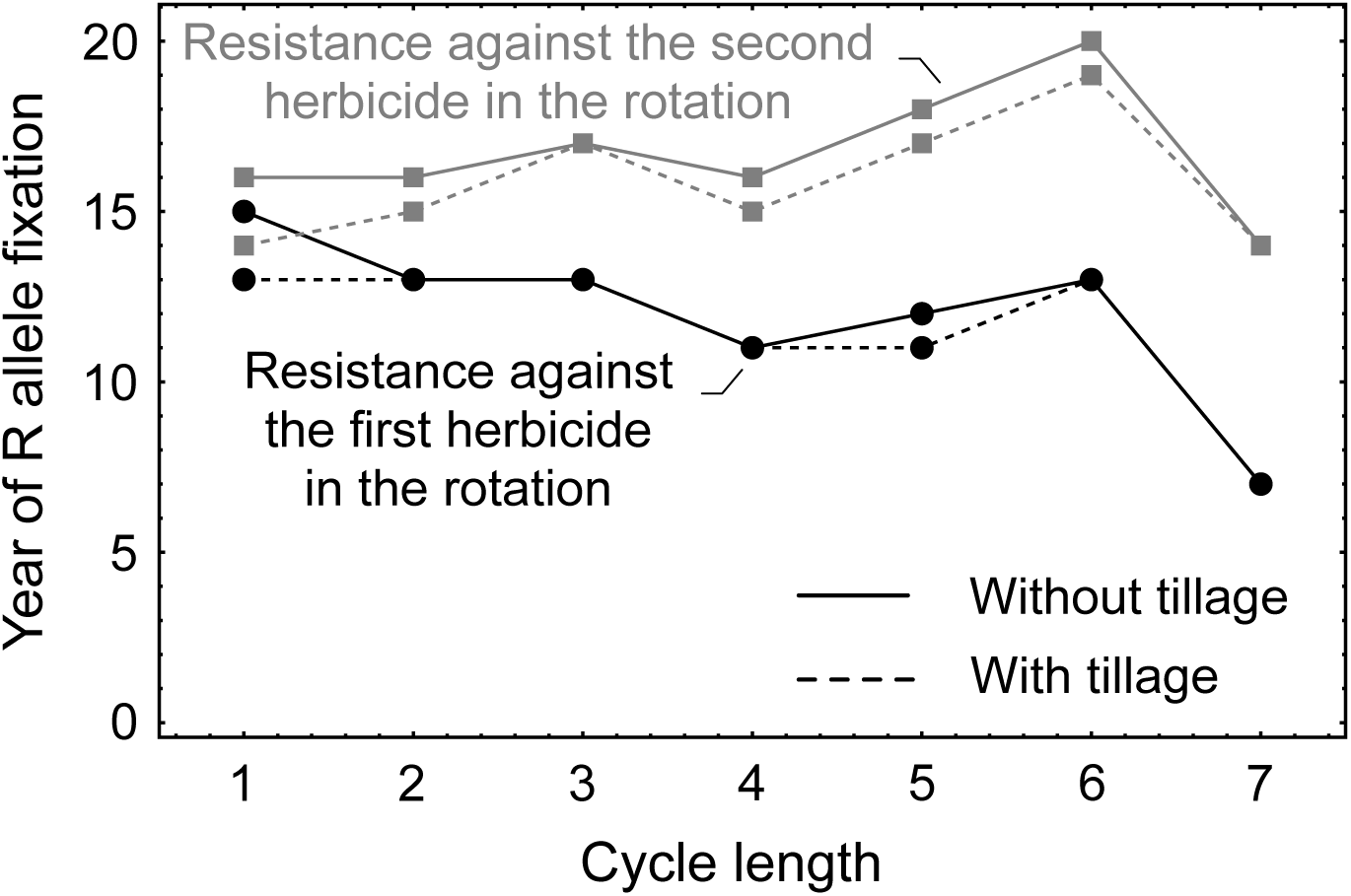
Predicted fixation time of target-site resistance in Johnsongrass under binary herbicide rotations. The results are obtained for a partially dominant resistance allele (*k_h_* = 0.5) and fitness cost (*k_c_* = 0.5). Considered are rotations of two herbicides with distinct target sites, ACCase- and ALS-inhibitors, Johnsongrass is known to develop target-site resistance towards. We assume equal efficacy here. The cycle length refers to the number of years one herbicide is recurrently applied before the treatment switches to the other herbicide. The year of resistance allele fixation is when the resistance allele frequency in plants (*P*), obtained from the deterministic model, reaches 99.5 %. The black lines with closed circles show the fixation of target-site resistance against the first herbicide in the rotation, and the grey line with closed rectangles resistance fixation against the second herbicide applied, where the herbicides are applied without mechanical control (solid line) or combined with tillage (dashed line).

